# Loss of MYB34 transcription factor that controls feedback loop in indole glucosinolate biosynthesis supports backward evolution of this pathway within Camelineae tribe

**DOI:** 10.1101/2022.06.29.496778

**Authors:** Paweł Czerniawski, Mariola Piślewska-Bednarek, Anna Piasecka, Karolina Kułak, Paweł Bednarek

**Author notes:** Corresponding author: Paweł Bednarek.

## Abstract

Glucosinolates are specialized defensive metabolites characteristic for the Brassicales order. Among them aliphatic and indolic glucosinolates (IGs) are usually highly abundant in the species from Brassicaceae family. The exception from this trend is constituted by species representing a subclade of Camelineae tribe, including *Capsella* and *Camelina* genera, which have reduced capacity to produce and metabolize IGs. Our study addresses contribution of specific glucosinolate-related MYB transcription factors to this unprecedented backward evolution of IG biosynthesis. To this end we performed phylogenomic and functional studies of respective MYB proteins. Obtained results revealed weakened conservation of glucosinolate-related MYB transcription factors, including loss of functional MYB34 protein, in the investigated species. We showed that introduction of functional MYB34 from *Arabidopsis thaliana* partially restores IG biosynthesis in *Capsella rubella* indicating that loss of this transcription factor contributes to the backward evolution of this metabolic pathway. Finally, we performed analysis of the impact of particular *myb* mutations on the feedback loop in IG biosynthesis, which drives auxin overproduction, metabolic dysregulation and strong growth retardation caused by mutations in IG biosynthetic genes. This uncovered unique function of MYB34 among IG-related MYBs in this feedback regulation and consequently in IG conservation in Brassicaceae plants.

## Introduction

Collectively, plants are capable to produce an enormous range of diverse specialized metabolites, which can play numerous functions in the interactions of plants with their environment. This huge chemical diversity is supported by continuous evolution of respective enzymes that can result in sub-branching of already existing metabolic pathways or in emergence of completely novel compound classes (Weng *et al*., 2012). In parallel, in particular species older pathways with diminished biological significance can undergo backward evolution that results in reduction of intraspecies chemical diversity. However, molecular mechanisms supporting such evets remain obscure.

One of the best recognized groups of plant specialized metabolites is constituted by glucosinolates, which are produced by species from the Brassicales order and include two major classes: aliphatic (AGs, mainly Met-derived) and indolic glucosinolates (IGs, Trp-derived) (Halkier and Gershenzon, 2006, Blažević *et al*., 2020). These sulfur-containing metabolites are produced constitutively through a conserved pathway involving precursor modification, core biosynthesis and glucosinolate modification (Sønderby *et al*., 2010b, Jensen *et al*., 2014, Barco and Clay, 2019). In the first step of core biosynthesis, chain elongated Met or Trp are converted into the corresponding aldoximes by cytochrome P450 monooxygenases CYP79, and then by monooxygenases CYP83 into *aci*-nitro compounds. Among these enzymes, CYP79B2, CYP79B3 and CYP83B1/SUR2 (Supperroot2) are specific for IG pathway (Fig. 1), while CYP79F1, CYP79F2 and CYP83A1 are involved in AG formation (Mikkelsen *et al*., 2000, Bak *et al*., 2001, Zhao *et al*., 2002, Chen *et al*., 2003, Naur *et al*., 2003). Subsequently, aliphatic and indolic intermediates are converted in several enzymatic steps, including action of S-alkyl-thiohydroximate lyase SUR1 and UDP-glucosyltransferases, to form glucosinolates (Grubb *et al*., 2004, Mikkelsen *et al*., 2004).

**Fig. 1.**
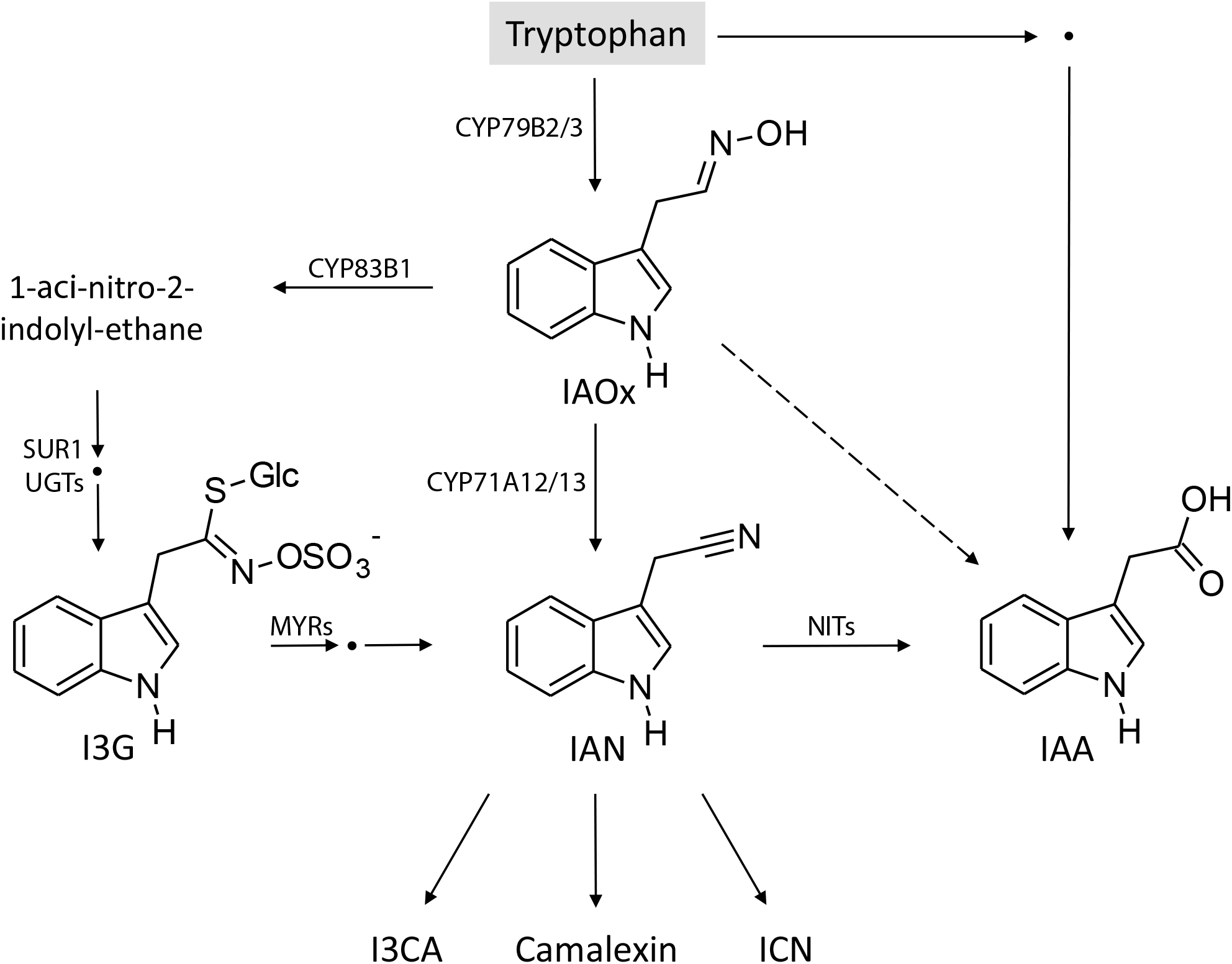
Simplified scheme of biosynthesis of indole glucosinolates and other Trp-derived metabolites. Only selected enzymes and intermediates are shown. SUR1 – Superrot1 C-S lyase, UGT – UDP-glycosyltransferase, MYR-myrosinase, NIT – nitrilase, IAOx – indole-3-acetaldoxime, IAN – indole-3-acetonitrile, IAA – indole-3-acetic acid, I3G – indolyl-3-methyl glucosinolate, ICN – indole-3-carbonyl nitrile, I3CA – indole-3-carboxylic acid. Multiple arrows represent multiple metabolic steps. Dashed arrow indicates hypothetical IAN-independent biosynthetic link between IAOx and IAA.

Published results suggest AGs and IGs accumulate constitutively in all plant organs of species belonging to the Brassicaceae family. An interesting exemption from this trend was found in species from *Capsella, Camelina, Neslia* and *Catolobus* genera, representing a subclade of Camelineae tribe, that do not accumulate glucosinolates in leaves (Fig. S1) (Windsor *et al*., 2005, Bednarek *et al*., 2011, Kiefer *et al*., 2014, Czerniawski *et al*., 2021). In other organs, these species accumulate predominantly AGs with relatively low levels of IGs. Moreover, the same species revealed reduced structural diversity of IGs and lost some of the genes encoding key enzymes required for function of IGs in plant immunity (Bednarek *et al*., 2011, Czerniawski *et al*., 2021). These findings suggested that IG biosynthetic capacity is on an evolutionary route to be shut down in this tribe. However, molecular mechanisms supporting this backward evolution of otherwise strongly conserved pathway remain unclear. Such mechanisms would be not only of interest in the context of glucosinolate pathway, but also regarding the evolution of specialized metabolites and chemical diversity in plants in general.

Biosynthesis of glucosinolates is tightly controlled by transcription factors belonging to the R2R3 MYB and basic helix-loop-helix (bHLH) MYC families (Frerigmann, 2016, Mitreiter and Gigolashvili, 2021). R2R3 MYB protein class is characterized by the presence of R2 and R3 domains, each containing three α-helices. The last helix of both domains coordinates the binding of the MYB protein to specific promoter sequences of the controlled genes (Kelemen *et al*., 2015). In *A. thaliana,* six MYBs have been reported as positive regulators of glucosinolate production and associated reactions including amino acid biosynthesis and sulfur assimilation (Frerigmann, 2016, Mitreiter and Gigolashvili, 2021). Previous studies have shown MYB28 and MYB29 play major role in biosynthesis of AGs in *A. thaliana,* with a minor contribution from MYB76 (Sønderby *et al*., 2007, Gigolashvili *et al*., 2008, Sønderby *et al*., 2010a). The strongest impact on regulation of IG biosynthesis has been reported for MYB34/ATR1 (Altered Tryptophan Regulation 1) and MYB51, while MYB122 has only a lower impact on the biosynthesis of these compounds (Celenza *et al*., 2005, Gigolashvili *et al*., 2007, Frerigmann and Gigolashvili, 2014). Among MYCs, function in glucosinolate biosynthesis in *A. thaliana* has been assigned to MYC2/bHLH06, MYC3/bHLH05/ATR2, MYC4/bHLH04, and MYC5/bHLH28 (Smolen *et al*., 2002, Dombrecht *et al*., 2007, Schweizer *et al*., 2013, Frerigmann *et al*., 2014). Of note, unlike MYBs, these proteins do not demonstrate specificity in the regulation of AG and IG production. As indicted by published results, proper control of glucosinolate biosynthesis requires direct interaction of MYB and MYC transcription factors mediated by MYC-interaction motif (MIM) locating downstream of the R3 domain (Frerigmann *et al*., 2014, Millard *et al*., 2019).

Tight control of glucosinolate biosynthesis seems to be essential as glucosinolates together with their activating *β*-thioglucosidases (myrosinases) play critical roles in defense of Brassicaceae plants against insect pests and microbial pathogens, as well as in response to abiotic stresses (Hopkins *et al*., 2009, Pastorczyk and Bednarek, 2016, Sugiyama *et al*., 2021). In addition, IG pathway is interconnected with the biosynthesis of other defensive Trp-derivatives and with the biosynthesis of plant hormone indole-3-acetic acid (auxin; IAA; Fig. 1) (Zhao *et al*., 2002, Bednarek, 2012, Frerigmann *et al*., 2016, Klein and Sattely, 2017, Malka and Cheng, 2017). The link between IG and IAA biosynthesis became obvious with isolation of *sur2/cyp83b1* mutant in a screen for mutants defective in auxin homeostasis. This mutant is characterized with elevated IAA levels, metabolic dysregulation, strongly inhibited shoot development and root extension at seedling stage, and severe growth retardation at later developmental stages (Delarue *et al*., 1998, Barlier *et al*., 2000, Bak and Feyereisen, 2001, Morant *et al*., 2010). As *cyp83b1* mutation leads to activation of Trp-biosynthesis and elevated expression of *CYP79B2/3* genes, it has been hypothesized that IG-deficiency activates initial step in biosynthesis of these metabolites resulting in elevated production of indole-3-acetaldoxime (IAOx), which in turn is channeled into IAA (Fig. 1) (Smolen and Bender, 2002, Naur *et al*., 2003, Celenza *et al*., 2005). Similar growth phenotypes and IAA overproduction have been reported in two other IG-deficient mutants, *sur1* and *ugt74b1* (Fig. 1) (Boerjan *et al*., 1995, Grubb *et al*., 2004, Mikkelsen *et al*., 2004), suggesting this mechanism is rather general for deficiency in IG biosynthetic enzymes acting downstream of IAOx. Moreover, this mechanism supports IG conservation handicapping individuals with spontaneous mutations in genes encoding respective enzymes, which could potentially lead to IG-deficiency. Apart from elevated expression of *CYP79B2/3* genes, *cyp83b1* seedlings demonstrated increased levels of *MYB34* transcript. Moreover, *myb34* mutation to some extent reverted expression of *CYP79B2/3* genes as well as growth phenotypes in *cyp83b1* seedlings indicating the mechanism inducing IAOx biosynthesis upon IG-deficiency is at least partially controlled by MYB34 (Celenza *et al*., 2005). However, potential impact of *myb51* and *myb122* mutations on this phenomenon was not investigated.

In this study, we addressed a hypothesis that evolutionary changes in transcription factors controlling IG biosynthesis contributed to backward evolution of this pathway in plants from clade II of Camelineae tribe (Fig. S1). Our phylogenomic analysis revealed weakened conservation of MYB transcription factors in this clade. Particularly we showed that loss of functional MYB34 has a significant impact on reduced production of IGs in these species. Moreover, we revealed unique role of MYB34 in the feedback loop controlling IG biosynthesis, which additionally highlights biological significance of loss of this particular transcription factor in backward evolution of glucosinolate pathway.

## Results

### *MYB34* is not fully conserved within the Camelineae tribe

We hypothesized that deficiency in IG production in species representing clade II of Camelineae tribe results from changes in activity of transcription factors controlling production of these compounds. Among transcription factors mediating glucosinolate biosynthesis, only MYB but not MYC proteins reveal specificity towards IG and AG production, hence, we focused on these transcription factors. We first looked for orthologs of *Arabidopsis thaliana* (L.) Heynh. *MYB* genes encoding proteins known to mediate glucosinolate biosynthesis in sequenced genomes of members of clade II of Camelineae tribe, including *Capsella rubella* Reut., *Capsella grandiflora* (Fauché & Chaub.) Boiss. and *Camelina sativa* (L.) Crantz (Fig. S1) (Slotte *et al*., 2013, Kagale *et al*., 2014). To have a broader insight into the conservation of these genes, we also included genomic sequences of *Arabidopsis lyrata* (L.) O’Kane & Al-Shehbaz (Hu *et al*., 2011) and of *Cardamine hirsuta* L., a species representing the Cardamineae tribe that is closely related to Camelineae (Gan *et al*., 2016). We found putative orthologs of *MYB28, MYB29, MYB51* and *MYB122* in genomes of all analyzed species (Fig. S2, Fig. S3). However, syntenic regions from all three copies of *Cam. sativa* genome did not contain *MYB34* orthologs (Fig. S2C). In addition, we did not find any *MYB76* orthologs in the analyzed genomes of species representing clade II of Camelineae tribe (Fig. S3B).

To further validate our ortholog identification, we performed phylogenic analysis of proteins encoded by these genes. We noted that the available sequences of MYB51 and MYB122 proteins from *C. grandiflora* were truncated at their N-termini (Fig. S4, Fig. S5). As the corresponding cDNA sequences deposited in the database did not include start codons we assumed this is an artifact resulting from the quality of *C. grandiflora* genome sequence, particularly that *MYB51* and *MYB122* genes are located at scaffold terminal regions (Fig. S2AB). To avoid any artifacts in the generated phylogenetic tree caused by such pseudo deletions, we used truncated sequences of all MYB proteins starting from Trp residue initiating α-helix 2 of R3 domain in our phylogenetic analysis (Fig. 2B, S4, S5). Resulting phylogenetic tree grouped respective orthologs into separate branches confirming our initial ortholog assignments (Fig. 2A). However, we noted that the relations within MYB28, MYB34 and MYB122 branches were incongruent with the phylogeny of species included in this analysis (Fig. 2A, Fig. S1). Proteins encoded by the identified *MYB29, MYB34* and *MYB122* orthologs from *A. thaliana* and *A. lyrata* showed closer relation to the respective enzymes from *Car. hirsuta,* which represents another tribe, than to those encoded by respective orthologs from remaining species representing Camelineae.

**Fig. 2.**
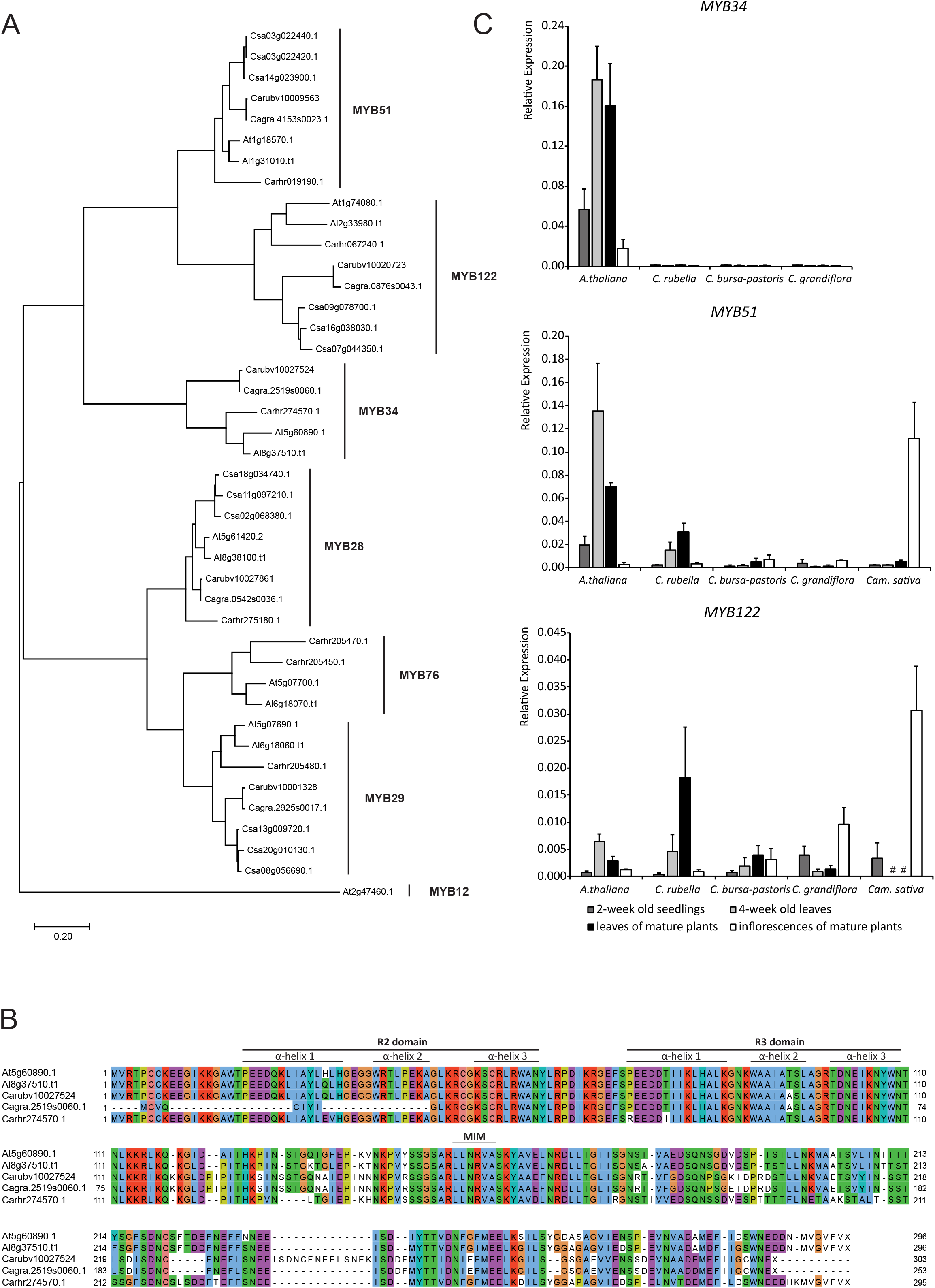
Comparative sequence and gene expression analysis of major MYBs known to regulate glucosinolate biosynthesis. (A) The maximum-likelihood phylogenetic tree of MYB proteins involved in regulation of IG biosynthesis from the investigated species. AtMYB12 was used as an outgroup. The bar represents 20% sequence divergence. (B) Alignment of proteins encoded by the identified MYB34 orthologs. R2, R3 and MIM (MYC-interaction motif) domains are indicated. (C) Relative expression levels of MYB34, MYB51, and MYB122 orthologs. Results are means ± SD from three independent experiments, each with four biological replicates (n = 12).

Due to our interest in IG biosynthesis, we checked MYB34 and MYB122 sequences and found that the observed phylogenetic incongruence correlated with several amino acid exchanges, deletions and insertions observed specifically in the sequences of proteins encoded by *MYB34* and *MYB122* orthologs from both analyzed *Capsella* spp. as well as in the three copies of *MYB122* found in *Cam. sativa* genome (Fig. 2B, S5). Particularly striking was the exchange of the conserved Thr to Ala in the α-helix 2 of R3 domain of MYB34 from *C. rubella* and *C. grandiflora,* as well as an insertion of 14 amino acids in the conserved region of C-terminal part of CrMYB34 (Fig. 2B). We also found an 11 amino acid deletion in the conserved sequence between R3 and MIM domains in both MYB122 proteins from *Capsella* spp. (Fig. S5). Moreover, we noted bigger deletions in the R2 domain of CgMYB34 and a deletion spanning R2 and R3 domains of two isoforms of CsMYB122 (Fig. 2B, Fig. S5). Collectively, all these alternations suggested that proteins encoded by *MYB34* and *MYB122* orthologs from species representing clade II of Camelineae tribe might have lost their function.

To support our ortholog analysis, we checked their expression levels in leaves and inflorescences of A. *thaliana, C. rubella, C. grandiflora, Capsella bursa-pastoris* (L.) Medik. and *Cam. sativa.* Our RT-qPCR analysis revealed that the relative transcript levels of *MYB28, MYB29, MYB34* and *MYB51* were usually lower in leaves of tested *Capsella* spp. and in *Cam. sativa* as compared with *A. thaliana* (Fig. 2C, Fig. S6). This was particularly striking for *MYB34* orthologs whose expression was in principle hardly detectable in all analyzed samples from tested *Capsella* spp. Overall, our results suggested that *MYB34* gene has been either lost or its expression is strongly compromised in species from clade II of Camelineae plants (Fig. 2C, Fig. S2).

### CrMYB34 is not functional in IG biosynthesis

Prompted by our results obtained during phylogenetic and gene expression analysis, we decided to test if MYB34 protein from *C. rubella* retained its function in IG biosynthesis. To this end, we expressed *CrMYB34* in *A. thaliana myb34 myb51 myb122* triple mutant that is deficient in IG biosynthesis. We assumed that functional *MYB34* ortholog should partially restore IG production in this mutant when expressed under the native *AtMYB34* promoter. We used RT-qPCR analysis with primers universal for *AtMYB34* and *CrMYB34* to screen T_1_ plants and to select two transgenic lines showing *CrMYB34* transcript at a similar level as *AtMYB34* transcript in Col-0 plants. These expression levels were further confirmed in T_2_ plants (Fig. 3A). We used the same lines to check with targeted LC-MS analysis the impact of *CrMYB34* expression on IG biosynthesis. This revealed no clear changes in accumulation levels of these compounds in leaves of the obtained transgenic lines as compared with the parental *myb34 myb51 myb122* plants (Fig. 3B). As expected, expression of *CrMYB34* did not affect AG accumulation (Fig. S7A). Collectively, these results indicated that *CrMYB34* lost its function in controlling IG biosynthesis.

**Fig. 3.**
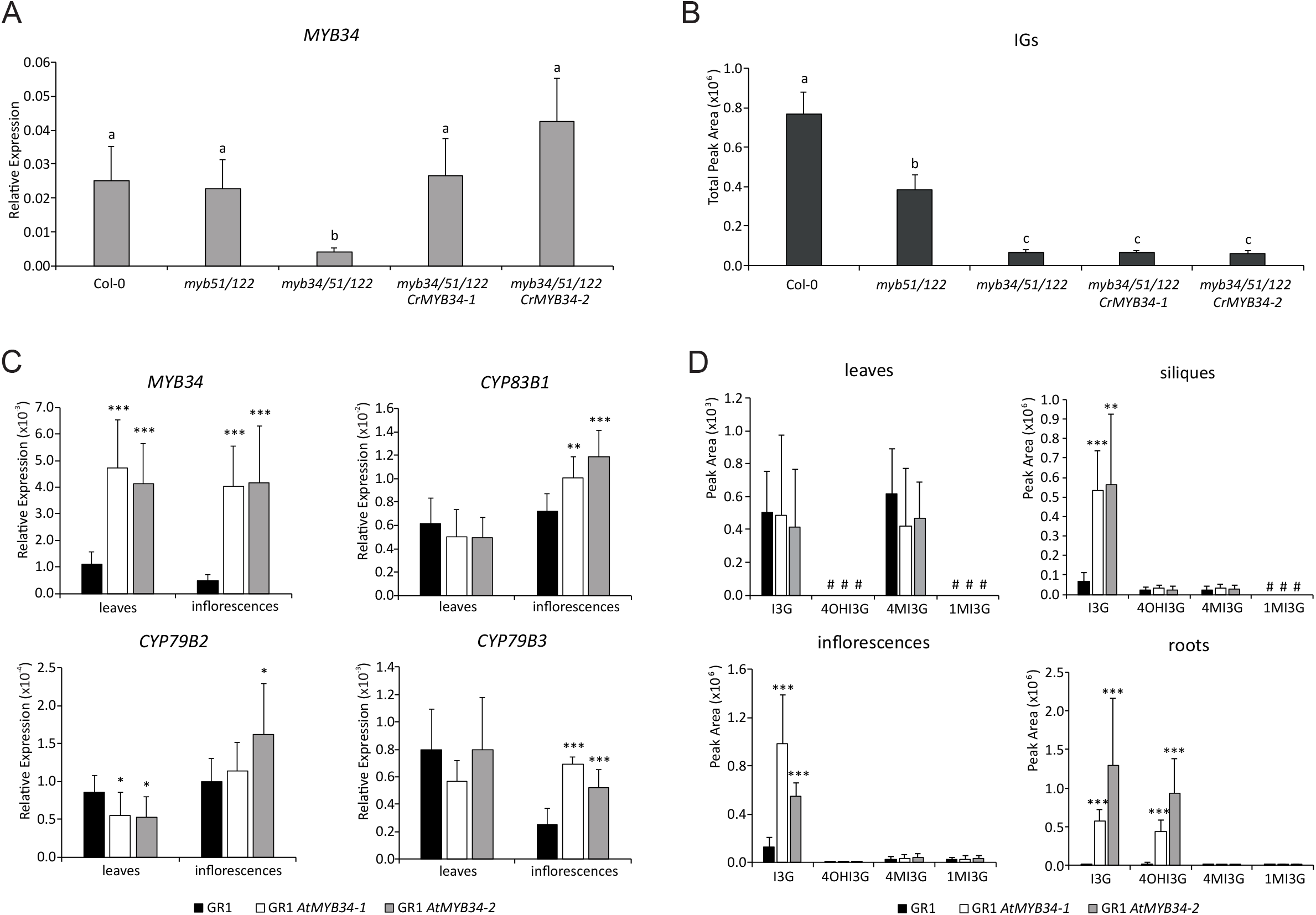
Loss of functional MYB34 contributes to indole glucosinolate (IG)-deficiency in *C. rubella*. (A) Relative expression of MYB34 gene in *A. thaliana myb* mutant and transgenic lines expressing *CrMYB34.* Results are means ± SD from five biological replicates (n = 5). Significantly different statistical groups are indicated based on ANOVA (P < 0.05, Games-Howell’s test). (B) Total peak areas of ions corresponding to IGs detected during LC-MS analysis of transgenic *A. thaliana* plants. The results are means ± SD from five biological replicates (n = 5). Significantly different statistical groups are indicated by ANOVA (P < 0.05, Tukey’s test). (C) Relative expression levels of selected genes regulated by MYB34 in transgenic *C. rubella* plants expressing *AtMYB34.*The results are means ± SD from three experiments, each with four biological replicates (n = 12). The values marked with asterisks are significantly different from respective controls (*, P < 0.05; **, P < 0.01; ***, P < 0.001). (D) Peak areas of ions corresponding to IGs identified in different organs of transgenic *C. rubella* lines expressing *AtMYB34.*The results are means ± SD from three experiments, each with four biological replicates (n = 12). The values marked with asterisks are significantly different from respective controls (**, P < 0.01; ***, P < 0.001). I3G – indolyl-3-methyl glucosinolate, 4OHI3G – 4-hydroxy-I3G, 4MI3G – 4-methoxy-I3G, 1MI3G – 1-methoxy-I3G, # - not detected.

### AtMYB34 partially restores IG biosynthesis in *C. rubella*

To check if the loss of functional MYB34 contributes to the reduced IG biosynthesis observed in the species from the clade II of Camelineae tribe, we expressed *MYB34* from *A. thaliana* under its native promoter in *C. rubella.* For practical reason, we decided to transform respective genetic construct into the GR1 ecotype (Slotte *et al*., 2006). Unlike the reference sequenced ecotype Monte Gargano, which was investigated in our earlier study and used for *MYB* expression analysis (Fig. 2C), flowering of GR1 does not require vernalization and consequently occurs faster. Before generating transgenic plants, we confirmed that the GR1 ecotype is similarly glucosinolate-deficient as Monte Gargano (Fig. S8) (Czerniawski *et al*., 2021).

We screened obtained T_1_ plants for those with elevated *MYB34* expression to select two of them and to confirm significantly higher levels of *MYB34* transcript in leaves and inflorescences of the respective T_2_ plants, as compared with wild-type GR1 plants (Fig. 3C). Subsequently, we checked if AtMYB34 can control expression of IG biosynthetic genes in *C. rubella.* To this end, we focused on the expression of previously identified orthologs of *A. thaliana* genes encoding CYP79B2, CYP79B3 and CYP83B1 monooxygenases specific for IG biosynthesis (Czerniawski *et al*., 2021). Respective RT-qPCR analysis revealed significantly higher expression levels of *CrCYP79B3* and *CrCYP83B1* in inflorescences of both tested lines expressing *AtMYB34* as compared with GR1 plants (Fig. 3C). Transcript level of *CrCYP79B2* was elevated slightly, but still significantly, only in one of the analyzed lines. Strikingly, despite detected expression of *AtMYB34* in leaves, we did not find any changes in the expression of analyzed genes in this organ (Fig. 3C). Overall, this indicted that AtMYB34 can at least partially control transcription of *CrCYP79B2, CrCYP79B3* and *CrCYP81B3.*

To check if the observed elevated expression of the analyzed genes results in enhanced IG biosynthesis, we investigated levels of these compounds in different organs of the obtained transgenic lines. This analysis revealed clearly higher accumulation of the unmodified indol-3-ylmethyl glucosinolate (I3G) in inflorescences, siliques and roots of both transgenic lines as compared with GR1 plants (Fig. 3D). In addition, roots of transgenic plants accumulated elevated levels of 4-hydroxy-I3G (4OHI3G). The amounts of the two remaining IGs, 1-methoxy-I3G (1MI3G) and 4MI3G were unaffected in the analyzed lines. As expected based on the gene expression profiles, IG accumulation in leaves was not affected with the expression of *AtMYB34* (Fig. 3CD). We also did not observe any impact of *AtMYB34* expression on the accumulation of AGs in the tested transgenic lines (Fig. S7B).

### MYB34 specifically controls overexpression of *CYP79B* genes in *A. thaliana cyp83b1* mutant

Our results revealed that functional MYB34 has been lost in the subclade II of Camelineae tribe, while at the same time MYB51 remained conserved in the same plant species. We wondered if there is any advantage in losing MYB34 over losing MYB51 during the backward evolution of IG biosynthesis. Interestingly, experimental results indicated MYB34 significantly contributes to the elevated *CYP79B2* and *CYP79B3* expression in *A. thaliana cyp83b1* seedlings (Fig. 1) (Celenza *et al*., 2005). However, the remaining IG-linked MYB transcription factors were not tested in this context, therefore it is not clear if this function is specific for MYB34, or shared with MYB51 and MYB122.

Concerning this, we decided to comprehensively test impact of *myb34, myb51* and *myb122* mutations on the activation of Trp-metabolism and the growth of *cyp83b1* mutant plants. The original *cyp83b1 myb34* double mutant has a mixed genetic background as it was generated by crossing *cyp83b1* mutant in Col-0 with *myb34 (atr1-2)* mutant in Ws background (Celenza *et al*., 2005). Additionally, we found that unlike reported *in vitro* under continuous light, *atr1-2* allele reveals strong growth retardation in soil under day/night cycle (Fig. S9). Therefore, we decided to generate our own uniform set of double *cyp83b1 myb* mutants by crossing isolated *cyp83b1* T-DNA line with *myb* mutants in Col-0 background.

We checked first generated double mutants for the expression levels of genes encoding MYB transcription factors and the three P450 monooxygenases specific for IG biosynthesis in leaves of 3-week old seedlings grown in soil. Our analysis revealed that *cyp83b1* mutation strongly upregulated expression of *MYB34,* but not of *MYB51* and *MYB122* (Fig. 4A, Fig. S10). As expected, we also found elevated *CYP79B2* and *CYP79B3* transcript levels in leaves of *cyp83b1* plants. This enhanced expression was completely reverted by *myb34* mutation. Despite their negative impact on *CYP79B2* and *CYP79B3* gene expression in the wild-type background, mutations in *MYB51* and *MYB122* did not significantly affect elevated expression of these two genes in the *cyp83b1* mutant (Fig. 3A). We also noted that lack of MYB34, but not of MYB51 or MYB122, reduced *CYP83B1* transcript level in wild-type background. We additionally found that transcript levels of *SUR1* that encodes C-S lyase contributing to the synthesis of IGs and AGs were unaffected in *cyp83b1* leaves (Fig. S10). Finally, we investigated expression of *CYP83A1,* which encodes P450 monooxygenase proposed to mediate IAOx conversion and IG biosynthesis in *cyp83b1* mutant (Naur *et al*., 2003). This revealed slightly reduced levels of *CYP83A1* transcript in leaves of *cyp83b1* mutant as compared with Col-0 (Fig. S10).

**Fig. 4.**
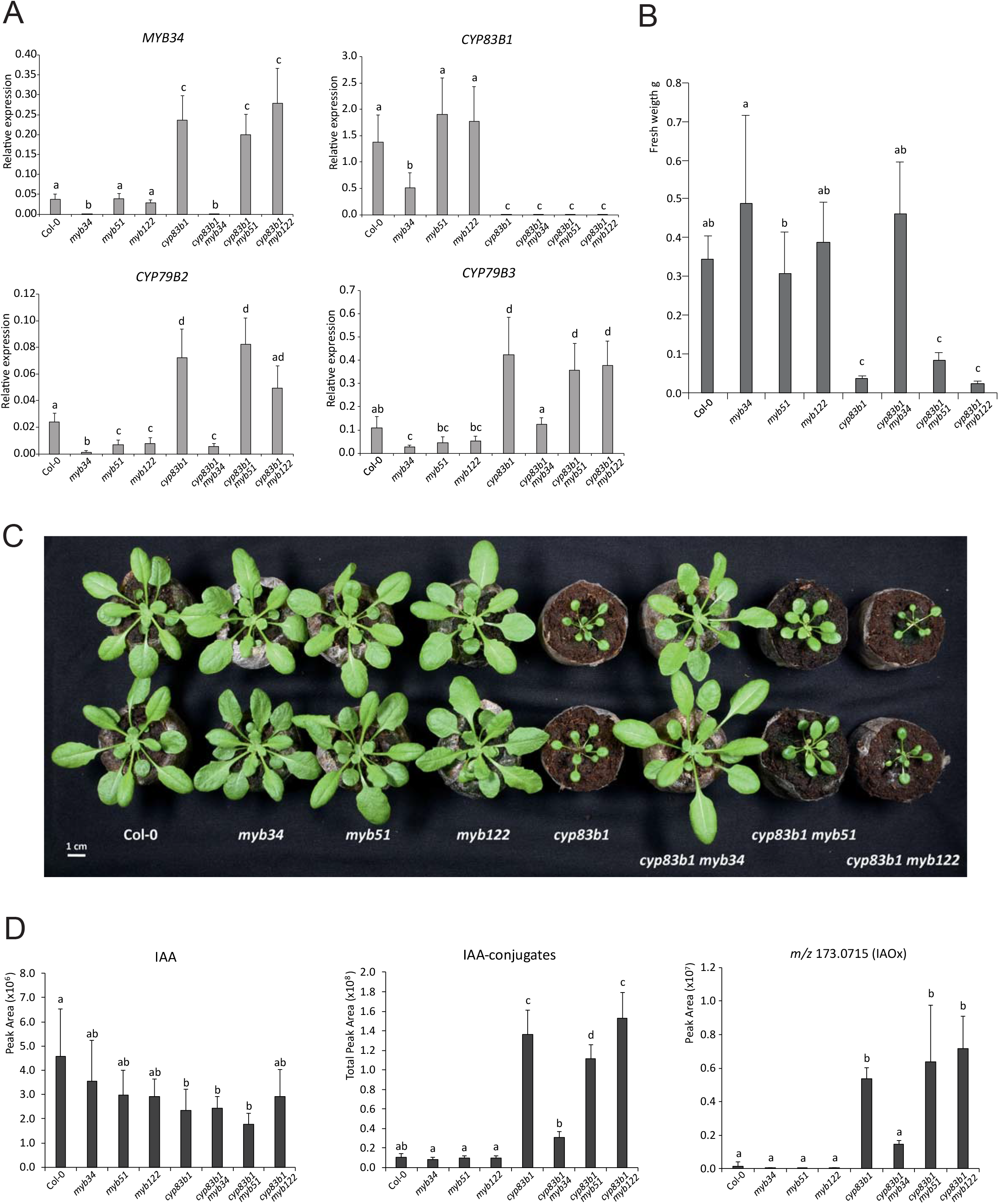
Impact of *myb* mutations on phenotypes of *A. thaliana cyp83b1* plants. (A) Relative expression levels of selected genes linked with indole glucosinolate biosynthesis in 3-week old leaves of the investigated *A. thaliana* mutant lines. Results are means ± SD from two experiments, each with four biological replicates (n = 8). Significantly different statistical groups are indicated based on ANOVA (P < 0.05, Games-Howell’s test). (B, C) Comparison of growth phenotypes of 6-week old *A. thaliana* mutant lines. Bar graph presents means ± SD from fresh weights of shoots collected in two experiments, each with eight biological replicates (n = 16). Significantly different statistical groups are indicated by ANOVA (P < 0.05, Tukey’s test). (D) Accumulation levels of indole-3-acetic acid (IAA), and of putatively identified IAA-derivatives and indole-3-acetaldoxime (IAOx) in leaves of 6-week old *A. thaliana* mutants. Bargraphs present peak areas corresponding to the respective ions detected during LC-MS analysis. Results are means ± SD from two experiments, each with four biological replicates (n = 8). Significantly different statistical groups are indicated based on ANOVA (P < 0.05, Tukey’s test).

### Mutation in *MYB34* reverts growth retardation and IAA hyperaccumulation in *cyp83b1* mutant background

To check if the investigated *myb* mutations can weaken growth retardation observed in mature *cyp83b1* plants, we grew generated double mutants together with Col-0 and respective single mutants for 6-weeks in soil and check fresh weight of plant rosettes. *cyp83b1* plants were strongly affected in their growth and fresh weight of their rosettes was about 10-times lower as compared with Col-0 (Fig. 4BC). *cyp83b1 myb51* and *cyp83b1 myb122* double mutants revealed similar growth phenotype. Unlike, *myb51* and *myb122, myb34* mutation turned back growth retardation caused by lack of functional CYP83B1 as indicated by fresh weight of *cyp83b1 myb34* rosettes, which was undistinguishable from the fresh weight of Col-0 rosettes (Fig 4BC).

The reduced growth of *cyp83b1* mutants has been proposed to be predominantly caused by hyperaccumulation of IAA, which is produced from the excess of IAOx. To check how mutations in particular *MYB* genes affect accumulation of these compounds, we performed targeted LC-MS analysis of samples isolated from leaves of 6-week old *cyp83b1 myb* mutants. First, we checked obtained data for *m/z* signal corresponding to IAOx [M-H]^-^ ion (Table S1). According to our analysis this signal was much more abundant in *cyp83b1* plants as compared with Col-0 (Fig. 4D). This was reverted by mutation in *MYB34,* but not in *MYB51* and *MYB122.* Next, we checked IAA accumulation by referring to the IAA standard. Oppose with the IAOx levels and unlike reported in seedlings (Delarue *et al*., 1998, Barlier *et al*., 2000), we found that free IAA levels were slightly, but significantly, reduced in leaves of *cyp83b1* mutant as compared with Col-0 plants (Fig. 4D). We assumed, this discrepancy could be caused by redirection of the excessive pools of IAA to respective derivatives. To test this hypothesis, we screened our LC-MS/MS data and found *m/z* signals corresponding to seven known IAA derivatives including conjugates with glucose and with six amino acids (Ala, Asp, Glu, Gly, Ser and Val) (Methods S1, Table S1). As indicated by the corresponding peak areas, unlike free IAA, all of these derivatives were present in significantly higher amounts in leaves of *cyp83b1* plants as compared with Col-0 (Fig. 4D, Fig. S11). Of note, *myb34,* but not *myb51* or *myb122,* mutation reverted this hyperaccumulation. Overall, our analysis indicated a clear correlation between growth retardation and enhanced accumulation of IAOx and IAA conjugates observed in leaves of *cyp83b1, cyp83b1 myb51* and *cyp83b1 myb122* plants.

### Majority of metabolic changes observed in *cyp83b1* plants is MYB34-dependent

As indicated by an earlier study, apart from IAA hyperaccumulation, metabolome of *cyp83b1* is clearly dysregulated as compared with Col-0 plants (Morant *et al*., 2010). Some of these metabolic changes result from accumulation of IAOx-derivatives other than IAA. We performed targeted HPLC-UV analysis to check accumulation of some of such compounds in leaves of our mutant set. As already reported, *cyp83b1* mutants counterintuitively are not IG-deficient due to the ability of CYP83A1 to convert IAOx when this intermediate appears at higher concentrations (Naur *et al*., 2003). Despite the slightly reduced *CYP83A1* expression levels in *cyp83b1* plants we did not note any clear IG deficiency in leaves of this mutant (Fig. 5A). We found only slight reduction in 4MI3G formation, which was compensated by elevated, as compared with Col-0 leaves, levels of 1MI3G. *myb34* mutation reduced accumulation of I3G and 4MI3G in the *cyp83b1* background to the levels observed in *myb34* single mutant, while *myb51* mutation in the *cyp83b1* background reduced only I3G accumulation. We checked also levels of glucoside of 6-hydroxy-indole-3-carboxylic acid (6OGIcI3CA) and I3CA glucose ester (I3CAGlc) (Fig. 1), and found profiles similar to those observed for IAOx and IAA-derivatives (Fig. 4D, Fig. 5A). Accumulation of these compounds was strongly enhanced in *cyp83b1* mutant and this was partially reverted by *myb34* mutation. Lack of MYB51 and MYB122 proteins did not affect amounts of both tested ICA derivatives (Fig. 5A). We also found weak, but statistically significant enhancement in accumulation of camalexin in *cyp83b1* plants (Fig. S11). To complete our investigation of known IAOx derived metabolites, we checked our LC-MS data for *m/z* signals representing indole-3-carbonylnitrile (ICN) and 4-hydroxy-ICN ions (Fig. 1) (Rajniak *et al*., 2015). Among those, we found only a signal potentially corresponding to ICN (Table S1). However, lack of CYP83B1 protein did not affect abundance of this ion in the respective samples suggesting that ICN formation in not affected by *cyp83b1* mutation (Fig. S11). Finally, we checked accumulation levels of indole-3-acetonitrile (IAN), which is one of the I3G hydrolysis products, can be also produced by CYP71A12/A13 enzymes on the route to ICA, ICN and camalexin, and has been proposed as an intermediate connecting IAOx and IAA (Fig. 1). However, we did not observe any changes in the accumulation of this compound in *cyp83b1* mutant as compared with wild-type plants (Fig. S11).

**Fig. 5.**
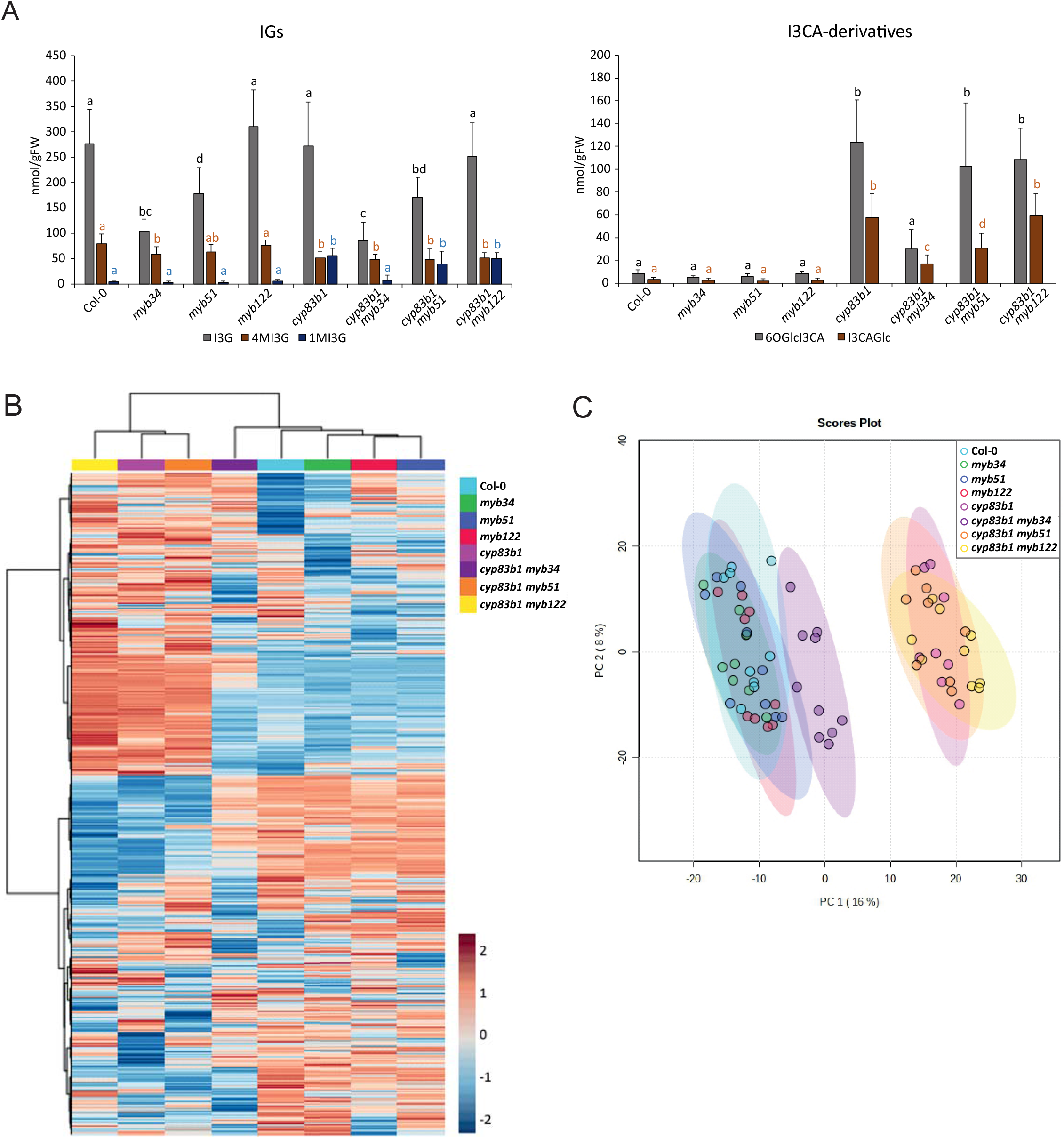
Metabolomic analysis of single and double *cyp83b1* and *myb A. thaliana* mutant lines. (A) Accumulation levels of indolic glucosinolates (IGs) and indole-3-carboxylic acid (I3CA)-derivatives in leaves of 6-week old *A. thaliana* mutants calculated based on HPLC-UV analysis. Results are means ± SD from two experiments, each with four biological replicates (n = 8). Significantly different statistical groups are indicated based on ANOVA (P < 0.05, Tukey’s test). I3G – indolyl-3-methyl glucosinolate, 4MI3G – 4-methoxy-I3G, 1MI3G – 1-methoxy-I3G, 6OGlcI3CA – I3CA 6-O-glucoside, I3CAGlc – I3CA glucose ester. (B) Hierarchical clustering of 2500 LC-MS signals significantly differentiating (one-way ANOVA; p < 0.01) leaf metabolomes of analysed lines. Heatmap presents peak areas of particular signals in log10 scale. (C) Score plot obtained for Principal Component Analysis (PCA) performed with the same set of 2500 signals.

To check the impact of *myb34* mutation on the accumulation of other compounds whose levels are also affected in *cyp83b1* plants, we performed untargeted processing of data obtained during our LC-MS analysis. Based on one-way ANOVA (p < 0.01), we selected 2500 signals significantly differentiating analyzed lines and performed their hierarchical clustering. Resulting dendrogram revealed two major sample clusters (Fig. 5B). One of those included *cyp81b1* together with *cyp83b1 myb51* and *cyp83b1 myb122* double mutant lines. The other included Col-0 together with single *myb* mutants and *cyp83b1 myb34* double mutant. To further validate these results, we conducted principal component analysis with the same set of signals. Similarly, as hierarchical clustering, this analysis grouped *cyp81b1, cyp83b1 myb51* and *cyp83b1 myb122* samples together. At the same time *cyp83b1 myb34* samples were strongly shifted towards, although did not overlap with, Col-0 and single *myb* mutant samples (Fig. 5C). Overall, our targeted and untargeted metabolomic analyses indicated that *myb34* mutation specifically reverts majority of metabolic changes caused directly or indirectly by *cyp83b1* mutation.

## Discussion

### Weakened conservation of glucosinolate-related MYBs in the Camelineae tribe

In this study, we investigated contribution of MYB transcription to the backward evolution of IG biosynthesis in a subclade of Camelineae tribe. Our phylogenomic analysis indicated that among the three MYBs controlling IG formation, only MYB51 is strongly conserved within the whole Camelineae tribe (Fig. 2A, Fig. S2A). Phylogenetic relationships between proteins encoded by *MYB34* orthologs from *C. rubella, C. grandiflora* and the remaining investigated species appeared to be inconsistent with species relatedness. Moreover, the hexaploid *Cam. sativa* appeared to be deprived from all three copies of *MYB34* ortholog suggesting that already the diploid ancestor of this species lost this gene (Fig. S2C). Similarly, as for the investigated MYB34 isoforms, we observed phylogenetic incongruence for proteins encoded by *MYB122* orthologs from the representatives of Camelineae clade II (Fig. 2A, Fig. S2B). Overall, this suggested that, unlike MYB51, MYB34 and MYB122 are no longer strongly conserved in species belonging to this clade.

The observed lack of straight conservation is not restricted to IG-related MYBs. MYB29 proteins do not follow phylogenetic relationships between investigated species suggesting that also this transcription factor is not strongly conserved within Camelineae tribe (Fig. 2A). Moreover, we did not find *MYB76* orthologs in analyzed genomes of species representing Camelineae subclade II (Fig. S3B). However, as MYB76 has only a weak impact on AG biosynthesis in *A. thaliana,* loss of this transcription factor seems to be without any stronger biological significance (Beekwilder *et al*., 2008, Gigolashvili *et al*., 2008, Sønderby *et al*., 2010a). Overall, among MYB proteins reported to control AG biosynthesis in A. *thaliana,* only MYB28 is strongly conserved in Camelineae tribe. As indicted by the analysis of respective *A. thaliana* mutant lines, MYB28 controls more specifically biosynthesis of long chain AGs while formation of short chain AGs is controlled jointly by MYB28 and MYB29 (Sønderby *et al*., 2007, Beekwilder *et al*., 2008). Concerning this, strong conservation of MYB28, but not of MYB29, in species from Camelineae clade II correlates with the reported capacity of these plants to produce predominantly long chain AGs (Czerniawski *et al*., 2021).

### Loss of functional MYB34 contributes to reduced IG biosynthesis in the subclade of Camelineae tribe

Mutations in *MYB34* genes from *Capsella* spp. combined with their marginal expression levels suggested these orthologs do not encode functional proteins (Fig. 2BC). Our experiments supported this hypothesis indicating that CrMYB34 is not able to even partially replace *At*MYB34 when expressed in *myb34 myb51 myb122* mutant under native *At*MYB34 promoter (Fig. 3AB). This likely results from particular exchanges of conserved amino acids, deletions or insertions observed in the sequence of CrMYB34. As CgMYB34 retains majority of these mutations and additionally possesses a deletion in the R2 domain (Fig. 2B), this protein likely is also no longer functional. These observations combined with lack of *MYB34* orthologs in *Cam. sativa* indicate that *MYB34* is on its way to be completely lost in the clade II of Camelineae tribe.

Conversely to the results obtained with CrMYB34, expression of functional *At*MYB34 under its native promoter in C. *rubella* partially restored IG production in the generated transgenic lines revealing that loss of this transcription factor significantly contributes to the reduced capacity of investigated Camelineae species to produce IGs (Fig. 3CD). However, despite elevated IG accumulation in GR1 *At*MYB34 lines, levels of these compounds were still lower than their accumulation in Col-0 plants indicating additional molecular determinants impairing IG biosynthetic capacity in this species (Fig. S7C). Possible candidates include MYB122, which is also not strongly conserved in Camelineae clade II (Fig. 2A, Fig. S5). However, as this transcription factor has only a marginal impact on IG biosynthesis in *A. thaliana* (Frerigmann and Gigolashvili, 2014), its loss should not drastically impact accumulation of these compounds.

Despite clear expression of *At*MYB34 in leaves of our transgenic *C. rubella* plants, we did not observe any increase in IG accumulation in this organ (Fig. 3CD) indicating other molecular features determining strong IG deficiency in leaves. As leaves of species from clade II of Camelineae tribe are IG- and AG-deficient (Czerniawski *et al*., 2021), it seems plausible that these factors simultaneously affect biosynthesis of all glucosinolates. Candidates for such molecular determinants include MYC-bHLH transcription factors, which do not reveal any specificity in controlling expression of genes encoding enzymes involved in IG and AG biosynthesis (Frerigmann, 2016, Mitreiter and Gigolashvili, 2021). In this context, it seems of interest that genes encoding respective MYC proteins have different organular expression pattern. For example, the *MYC2* gene in *A. thaliana* is most highly expressed in roots, while the expression of *MYC3* and *MYC4* is the highest in shoot tissue and *MYC5* in leaves and inflorescences (Figueroa and Browse, 2015, Gasperini *et al*., 2015, Qi *et al*., 2015). Alternatively, lack of yet uncharacterized regulators of glucosinolate biosynthesis could be responsible for glucosinolate deficiency in leaves of investigated species.

Among all IGs, only I3G increased uniformly its accumulation levels in inflorescences, siliques and roots of *C. rubella* transgenic lines (Fig. 3D). This is in accordance with the findings that MYB transcription factors control core I3G biosynthesis, but not subsequent modification of this compound (Frerigmann and Gigolashvili, 2014). However, in addition to the increase of I3G concentration, we also found elevated levels of 4OHI3G in roots of the generated transgenic lines. This suggests *At*MYB34 might have direct or indirect impact on the formation of this compound in *C. rubella* root. This species lacks CYP81F2 monooxygenase that in *A. thaliana* is a key player in 4OHI3G formation (Bednarek *et al*., 2009, Bednarek *et al*., 2011). Alternative enzymes include CYP81F1 and CYP81F3, which were characterized in *A. thaliana* or CYP81F5 and CYP81F6 that are specific for Camelineae clade II, but their function in I3G modification has not been tested so far (Pfalz *et al*., 2011, Czerniawski *et al*., 2021). Finally, the strong increase in 4OHI3G levels in roots did not result in elevated 4MI3G production (Fig. 3D). This may support our earlier hypothesis that the respective O-methyltransferases lost their function in the clade II of Camelineae tribe (Czerniawski *et al*., 2021).

### MYB34 possesses a unique function in controlling IG levels

According with the published results, MYB34 and MYB51 are the key players among the three MYBs controlling IG biosynthesis (Frerigmann, 2016, Mitreiter and Gigolashvili, 2021). These transcription factors differ slightly in their tissue specificity with MYB34 being more active in roots and MYB51 in leaves. In addition, they respond differentially to particular plant hormones and environmental cues (Gigolashvili *et al*., 2007, Frerigmann and Gigolashvili, 2014, Frerigmann *et al*., 2016). Our study highlights another difference between transcription factors controlling IG biosynthesis. Results obtained during analysis of the series of *cyp83b1 myb* mutants indicated that MYB34, but not MYB51 or MYB122, specifically controls induced expression of *CYP79B2/3* genes caused by lack of CYP83B1 enzyme (Fig. 4A). Overall, these results suggest MYB34 acts specifically in a feedback loop activating expression of genes encoding pathway entry enzymes in response to IG deficiency. This in turn implies existence of molecular mechanisms negatively correlating expression of *MYB34* with IG accumulation levels, however, the details of this mechanism remain obscure. As metabolites can control gene expression by interaction with enzymes involved in chromatin modification, with proteins binding to gene promoter regions or with proteins involved in mRNA maturation (Ladurner, 2006), direct involvement of IGs in this regulation can be not excluded.

### Significance of MYB34 in conservation of IG biosynthesis

Overproduction of IAOx, and its derivatives is not the only metabolic effect of *cyp83b1* mutation. As indicated by earlier metabolomic studies, this mutation leads to massive metabolome dysregulation suggesting that despite from being metabolized to IAA and other Trp-derivatives, IAOx also impacts directly or indirectly other metabolic pathways (Morant *et al*., 2010). For instance, IAOx, or its derivative, has a negative impact on the expression of *Phenylalanine ammonia lyase* affecting phenylpropanoid accumulation in *cyp83b1* plants (Kim *et al*., 2015). All these metabolic changes contribute to growth retardation and eventually lethality observed not only in *cyp83b1,* but also in *sur1* and *ugt74b1* mutants (Boerjan *et al*., 1995, Delarue *et al*., 1998, Grubb *et al*., 2004). This in turn protects from evolutionary losses of IG biosynthetic capacity caused by spontaneous mutations in respective biosynthetic genes. Our results indicate that lack of MYB34, but not of MYB51 or MYB122, weakens the feedback loop and prevents from IAA overproduction, metabolic dysregulation and growth penalty (Fig. 4, Fig. 5). Consequently, loss of this particular MYB transcription factor may additionally support backward evolution of IG biosynthesis in clade II of Camelineae plants.

### Metabolic fate of IAOx excess and IAA formation in *cyp83b1* plants

Earlier studies revealed *cyp83b1* plants to be counterintuitively only partially IG-deficient. This has been explained with contribution of CYP83A1, which according to *in vitro* experiments is more specific towards aliphatic oximes, but still capable to efficiently convert IAOx at higher concentrations of this substrate (Bak and Feyereisen, 2001, Naur *et al*., 2003). Our study revealed maintained IG levels in *cyp83b1* leaves indicating that CYP83A1 can fully replace CYP83B1 at the observed elevated concentrations of IAOx (Fig. 4D, Fig. 5A). However, a study on glucosinolate biosynthetic enzymes fused with fluorescent proteins revealed that AG and IG biosynthesis localize to different cells. Consequently, under normal physiological conditions IAOx is produced in cells that do not express *CYP83A1* (Nintemann *et al*., 2018). This combined with our result suggests this spatial separation of both glucosinolate pathways is no longer maintained in *cyp83b1* mutants. Possibly, the MYB34-triggered increase in *CYP79B2/3* expression occurs not only in IG, but also in AG producing cells. However, other scenarios including IAOx transport/diffusion or aberrant cellular expression pattern of *CYP83A1* cannot be excluded.

Our analysis indicated that the excess of IAOx is preferentially redirected into IAA and I3CA biosynthesis (Fig. 4D, Fig. 5A, Fig. S11). Despite the shared with I3CA biosynthetic step catalyzed by CYP71A12 monooxygenase (Fig. 1) (Rajniak *et al*., 2015, Pastorczyk *et al*., 2020), we were not able to detect any increase in the amounts of ICNs in *cyp83b1* (Fig. S11). Conversely, we noted significant, but rather weak, increase in camalexin accumulation (Fig. S11). As both ICN and camalexin are toxic, it seems likely that these branches of Trp-metabolism are controlled more tightly than I3CA and IAA biosynthesis.

Although the biosynthetic link between IAOx and IAA has been confirmed with *cyp83b1* and *cyp79b2 cyp79b3* mutants, the details of this metabolic connection remain obscure (Barlier *et al*., 2000, Zhao *et al*., 2002). Published results indicate IAA can be formed by specific nitrilases from IAN, which is produced upon I3G hydrolysis or formed from IAOx by CYP71A12/13 monooxygenases (Fig. 1) (Normanly *et al*., 1997, Mano and Nemoto, 2012, Lehmann *et al*., 2017, Malka and Cheng, 2017). The first possibility is unlikely to explain IAA overproduction in *cyp83b1* plants as this would require elevated production and hydrolysis of I3G. Consequently, involvement of CYP71A12/13 enzymes seems to be more likely. However, in this case a correlation between concentrations of IAOx, IAN and IAA derivatives in *cyp83b1* plants would be expected and this was not confirmed with our analysis of IAN accumulation in leaves (Fig. S11). Of note, as indicated by *in vitro* studies CYP71A12/13 monooxygenases can convert IAOx to oxidized forms of IAN, rather than to IAN itself (Klein *et al*., 2013). It remains to be tested if such IAN derivatives could also be converted into IAA. Finally, other, yet uncharacterized, metabolic routes can also contribute to IAA overproduction in *cyp83b1* plants.

## Experimental procedures

### Plant material

*Capsella rubella* accession Monte Gargano and *Arabidopsis thaliana cyp83b1* mutant (SALK_071430) seeds were obtained from the Nottingham Arabidopsis Stock Centre (stock #N9609 and #N571430, respectively). *C. rubella* GR1 seeds were received from Prof. Tanja Slotte (Stockholm University, Sweden) (Slotte *et al*., 2006). *Capsella bursa-pastoris* seeds were from B & T World Seeds (stock #1507), *Capsella grandiflora* was obtained from Prof. Miltos Tsiantis (Max Planck Institute for Plant Breeding Research, Germany). *Camelina sativa* homozygous doubled haploid DH55 line was shared by Dr. Isobel Parkin (Saskatoon Research and Development Centre, Agriculture and Agri-Food Canada) (Kagale *et al*., 2014). The seeds of A. *thaliana myb34, myb51,* and *myb122* single and multiple mutants were obtained from Dr. Henning Frerigmann (University of Cologne, Germany) (Gigolashvili *et al*., 2007, Frerigmann and Gigolashvili, 2014). *cyp83b1 myb* mutant lines were generated by genetic crosses followed by selection of homozygous mutant alleles (Table S2).

### Generation of transgenic plants

We used pAMPAT binary vector (AY436765) to express genes of interest in A. *thaliana* and *C. rubella.* To express *AtMYB34,* we cloned a PCR amplified genomic fragment (A. *thaliana* Col-0) containing this gene together with its promoter region (1318 bp upstream from *AtMYB34* start codon) (Table S2). The same promoter region was also cloned into the pAMPAT vector in fusion with PCR amplified *CrMYB34 (C. rubella* Monte Gargano). The obtained plasmids were transformed into *Agrobacterium tumefaciens* strain GV3101:pMP90RK (Koncz and Schell, 1986), which was used for stable transformation of *C. rubella* GR1 and A. *thaliana myb34 myb51 myb122* plants with floral dip method. Transgenic *C. rubella* plants were selected with glufosinate (75 mg/l; Merck, USA) spray. As *myb34 myb51 myb122* plants retain partial glufosinate resistance linked with *myb34* (WiscDsLox424F3) and *myb122* (WiscDsLoxHs206_04H) T-DNA insertions, higher herbicide concentration (150 mg/l) was used in this case. Leaf samples were collected from resistant 6 weeks old T_1_ plants for RT-qPCR analysis to select plants with the highest transgene expression levels.

### Experimental conditions and sample collection

Camelineae species used for *MYB* gene expression analysis as well as wild-type and transgenic T_2_ *C. rubella* GR1 plants used for gene expression and glucosinolate analysis were grown and samples were collected as described earlier (Czerniawski *et al*., 2021).

A. *thaliana* Col-0, *myb* and *cyp83b1* mutants as well as T_2_ transgenic plants were grown in Jiffy-7 pellets (Jiffy Group, Netherlands) under short-day conditions (8 h light, 16 h darkness). Samples of 3-week old *cyp83b1* and *myb* mutants were collected for gene expression analysis. Leaf material collected from 6-week old plants was used for metabolomic analysis of the same mutants and for gene expression and metabolomic analysis of transgenic A. *thaliana* lines expressing *CrMYB34* gene.

### Ortholog identification and gene expression analysis

Gene and protein sequences were collected from genomic databases available for tested species as described earlier by Czerniawski *et al.* (2021). Protein sequences were aligned using MUSCLE algorithm and phylogenetic tree was constructed based on the maximum-likelihood algorithm in MEGA v10.1.7 software. Total RNA isolation from 50 mg frozen tissue, reverse transcription and RT-qPCR analysis were performed as reported earlier (Czerniawski *et al*., 2021). We designed single primer pairs matching all of the three putative orthologs of each gene identified in *Cam. sativa* allowing eventually single nucleotide mismatches. We designed common primers for *C. rubella* and *C. grandiflora* and used these primers also in *C. bursa-pastoris.* Similarly, we designed common primers for *AtMYB34* and *CrMYB34* to investigate MYB34 transgenic lines (Table S2).

### Metabolite extraction and analysis

Frozen plant samples (~200 mg) were homogenized in DMSO (2.5 μl per 1 mg fresh weight), centrifuged, and the supernatants were collected for further analysis.

#### Targeted LC-MS glucosinolate analysis

Glucosinolate detection and quantification was performed using ultra-performance liquid chromatography-tandem mass spectrometry (UPLC-MS/MS) system consisting of Acquity UPLC (Waters, USA) coupled with a micrOTOF-Q mass spectrometer (Bruker Daltonics, Germany) as described earlier (Czerniawski *et al*., 2021).

#### Targeted analysis of selected Trp-derivatives and untargeted metabolomic analysis

Targeted analysis of IAOx, indole-3-carbonyl nitrile, IAA and its conjugates, as well as untargeted analysis of leaf samples from 6-week old *A. thaliana* mutant lines, were conducted on Acquity UPLC (Waters, USA) hyphenated to Q-Exactive hybrid quadrupole Orbitrap mass spectrometer (Thermo Scientific, USA) (Table S3, Methods S1, Methods S2).

#### LC-UV-FLD analysis

Identification and quantification of IGs, indole-3-carboxylic acid (I3CA) derivatives, camalexin and indole-3-acetonitrile (IAN) were performed using Agilent 1200 HPLC System (Agilent, USA) equipped with diode array and fluorescence (FLD) detectors according to the published protocol (Bednarek *et al*., 2009, Pastorczyk *et al*., 2020). IAN was detected with FLD detector (excitation 275 nm, emission 350 nm). The amounts of selected metabolites in fresh tissue were calculated based on calibration curves obtained for standard compounds.

## Acknowledgements

This work was supported by the National Science Centre OPUS grant (2015/17/B/NZ1/00871) to PB. We would like to acknowledge Prof. Miltos Tsiantis (Max Planck Institute for Plant Breeding Research, Germany), Dr. Isobel Parkin (Saskatoon Research and Development Centre, Agriculture and Agri-Food Canada), Prof. Tanja Slotte (Stockholm University, Sweden) and (Dr. Henning Frerigmann (University of Cologne, Germany) for sharing the seeds. We also would like to thank Dr. Marta Pastorczyk-Szlenkier for assistance with crosses of *cyp83b1* and *myb* lines, and Dr. Gopal Singh for his comments on the manuscript.

## Author Contributions

PC, MPB and PB conceived the project and designed experiments. PC, MPB, AP and KK performed the experiments and processed the data with the supervision of PB. PC and PB wrote the manuscript with contributions from the remaining authors.

## Supporting Information

Additional Supporting Information may be found in the online version of this article.

---

**Fig. S1.** Phylogenetic tree representing relationships between species belonging to the Camelineae tribe.

**Fig. S2.** Graphical representation of genomic regions containing *MYB51, MYB122,* and *MYB34* orthologs in the investigated Camelineae species.

**Fig. S3.** Graphical representation of genomic regions containing *MYB28* and *MYB29/MYB76* orthologs in the investigated Camelineae species.

**Fig. S4.** Alignment of proteins encoded by *MYB51* orthologs identified in the investigated Camelineae species.

**Fig. S5.** Alignment of proteins encoded by *MYB122* orthologs identified in the investigated Camelineae species.

**Fig. S6.** Relative expression levels of *MYB28* and *MYB29* orthologs in the investigated Camelineae plants.

**Fig. S7.** Total accumulation of glucosinolates.

**Fig. S8.** Total accumulation of aliphatic and indolic glucosinolates in different organs and developmental stages of *C. rubella* GR1 ecotype.

**Fig. S9.** Comparison of growth phenotypes of 6-weeks old Col-0, Ws, *myb34,* and *atr1-2* plants.

**Fig. S10.** Relative expression levels of *MYB51, MYB122, CYP83A1,* and *SUR1* genes in *A. thaliana* single and double *cyp83b1* and *myb* mutant lines.

**Fig. S11.** Impact of *cyp83b1* and *myb* mutations on accumulation of different Trp-derivatives in *A. thaliana* leaves.

**Table S1.** Identification of Trp-derivatives with LC-MS/MS analysis.

**Table S2.** Sequences of primers used in this study.

**Table S3.** Heights and m/z values of peaks detected during LC-MS analysis of samples obtained from *cyp83b1 myb* mutant lines.

**Methods S1.** Targeted analysis of selected Trp-derivatives and untargeted metabolomic analysis.

**Methods S2.** Parameters of raw MS data processing by MZmine software.

## Notes

### Competing Interest Statement

The authors have declared no competing interest.

