## Supplementary files for "Loss of MYB34 transcription factor that controls feedback loop in indole glucosinolate biosynthesis supports backward evolution of this pathway within Camelineae tribe"

**Methods S1** Targeted analysis of selected Trp-derivatives and untargeted metabolomic analysis.

***Targeted analysis of selected Trp-derivatives***

Targeted analysis of IAOx, ICN, IAA and its conjugates, as well as untargeted analysis of 6-week old plant leaves of *A. thaliana* mutant lines, were conducted on Acquity UPLC (Waters, USA) hyphenated to Q-Exactive hybrid quadrupole Orbitrap mass spectrometer (Thermo Scientific, USA). Chromatographic separation was done on Acquity UPLC HSS T3 C18 column (2.1 x 50 mm, 1.8 µm particle size; Waters, USA) at 22°C. The elution was conducted with water containing 0.1% formic acid (solvent A) and acetonitrile (solvent B) at flow rate of 0.3 ml/min. The gradient elution was started at 98 % of A and linearly changed to 65 % of A over 4 min, then with two minutes changed to 2 % of A and was constant over 8 min. Q-Exactive MS operated by Xcalibur version 3.0.63 software with the following settings: heated electrospray ionization ion source voltage −3 kV; sheath gas flow 30 l/min; auxiliary gas flow 13 l/min; ion source capillary temperature 250°C; auxiliary gas heater temperature 380°C. MS/MS mode (data-dependent acquisition) was recorded in negative ionization, at resolution 70000 and AGC (ion population) target 3e6, with scan range 75 to 1000 m/z.

In order to identify IAA, we performed analysis of the standard of this compound. Fragmentation of IAA in negative ionization mode, leads to characteristic product ions at *m/z* 116.0492 and 130.0649 corresponding to $[C_8H_6N]^-$ and $[C_9H_8N]^-$ ions, respectively. In the case of fragmentation spectra of IAA conjugates with sugars and amino acids, apart from the above-mentioned product ions, signal at *m/z* 174. 0555 corresponding to the deprotonated IAA molecule are also observed. The *m/z* values of the molecular ions of the IAA conjugates were then compared with the available literature and data bases to identify particular IAA-conjugates (Tam *et al.*, 2000; Revelou *et al.*, 2019). In the case of IAOx and ICN, we did not obtain their fragmentation spectra, so we conducted further analyzes based on the signals at *m/z* 173.0715 and 169.0399 respectively, presumably corresponding to these deprotonated molecules. The level of accumulation of individual compounds was determined as peak areas on respective ion chromatograms (Table S2).

***Untargeted metabolomic analysis***

Raw LC-MS data obtained with Orbitrap mass spectrometer was processed for peak detection, deisotoping, alignment and gap filling by MZmine 2.51 (Methods S2) (Pluskal *et al.*, 2010). Processed data table contained 56857 signals, 8 groups with 9 biological repetitions  was imported to “Statistical Analysis [one factor]” module in MetaboAnalyst 5.0 for PCA and heatmap creation (Table S3) (Pang *et al.*, 2021). Imported data table was first filtered by interquartile range, normalized by $\log_{10}$ transformation and subjected to ANOVA with FDR correction for selection of statistically significant signals with p-value < 0.01. Selected 2500 signals were organized by decreasing p-value and clustered by WARD algorithm with similarity measurement on the basis of Euclidean distance. Score Plot for PCA was created on the same data with 95% confidence ellipse.

**Table S1** Identification of Trp-derivatives with LC-MS/MS analysis * - fragmentation ions not detected.

| No. | Retention time [min] | Metabolite | Chemical formula | Exact mass of [M-H]- | | error [ppm] | Fragmentation |
|---|---|---|---|---|---|---|---|
| | | | | Measured | Calculated | | |
| 1 | 2.97 | IAOx (indole-3-acetaldoxime) | $C_{10}H_{10}N_2O$ | 173.0715 | 173.0712 | -1.67 | * |
| 2 | 3.01 | IAGly (indole-3-acetyl-L-glycine) | $C_{12}H_{11}N_2O_3$ | 231.0770 | 231.0776 | 2.60 | 231, 174, 161, 130, 116 |
| 3 | 3.36 | IASer (indole-3-acetyl-L-serine) | $C_{13}H_{13}N_2O_4$ | 261.0875 | 261.0881 | 2.30 | 261, 174, 156, 130, 116, 104 |
| 4 | 3.39 | ICN (indole-3-carbonyl nitrile) | $C_{10}H_6N_2O$ | 169.0399 | 169.0402 | 1.46 | * |
| 5 | 3.61 | IAGlc (indole-3-acetyl-D-glucose) | $C_{16}H_{18}NO_7$ | 336.1083 | 336.1089 | 1.79 | 336, 280, 262, 174, 163, 130, 116 |
| 6 | 3.82 | IAAsp (indole-3-acetyl-L-aspartic acid) | $C_{14}H_{13}N_2O_5$ | 289.0824 | 289.0832 | 2.77 | 289, 262, 228, 161, 146, 128, 116, 102 |
| 7 | 3.94 | IAGlu (indole-3-acetyl-L-glutamic acid) | $C_{15}H_{15}N_2O_5$ | 303.0981 | 303.0988 | 2.31 | 303, 280, 221, 174, 146, 130 |
| 8 | 4.18 | IAAla (indole-3-acetyl-L-alanine) | $C_{13}H_{14}N_2O_3$ | 246.0926 | 246.0934 | 3.26 | 246, 228, 174, 160, 144, 130, 116 |
| 9 | 4.81 | IAA (indole-3-acetic acid) | $C_{10}H_8NO_2$ | 174.0555 | 174.0553 | -1.79 | 174, 159, 147, 130, 116, 100 |
| 10 | 5.22 | IAVal (indole-3-acetyl-L-valine) | $C_{15}H_{17}N_2O_3$ | 273.1239 | 273.1247 | 2.93 | 273, 174, 164, 130, 116 |

**Table S2** Sequences of primers used in this study. * primers used in the analysis of *MYB* gene expression in Camelineae species. # primers used in analysis of gene expression in *cyp83b1 myb* lines.

| | | |
|---|---|---|
| **A. Gene expression analysis by RT-qPCR.** | | |
| ***A. thaliana*** | | |
| At2g37620.1 | At_Actin_F | CCGGTATTGTGCTGGATTCT |
| | At_Actin_R | AATTTCCCGCTCTGCTGTTG |
| At4g39950.1 | At_CYP79B2_F | AACAAAAAGAAACCGTATCTGCCAC |
| | At_CYP79B2_R | TCCTAACTTCACGCATGCTATCTC |
| At2g22330.1 | At_CYP79B3_F | ACCGCTGATGAAATCAAACC |
| | At_CYP79B3_R | TCCGACGACTCTATCGATCT |
| At4g13770.1 | At_CYP83A1_F | CACATCGTGGCCATGAGTT |
| | At_CYP83A1_R | ACTTCGGATTTATCCGCGG |
| At4g31500.1 | At_CYP83B1_F | GAAATCTCTCCGGTTACCTC |
| | At_CYP83B1_R | TAGCTCGGCCGAGGAGATCA |
| At1g74080.1 | At_MYB122_F | GTTGTAGAGCAGAAGGGTTGA *<br>CGACGTCGTTTAATGATAGC # |
| | At_MYB122_R | TCTCAATCTGCAGCTTTTGC*<br>CTAACAACGTCAAAATTCACAA # |
| At5g61420.2 | At_MYB28_F | ACGTTTGATGGAACAGGGTATT |
| | At_MYB28_R | CCATCATTGCTGCTAAGCTC |
| At5g07690.1 | At_MYB29_F | GCTCCTGATCGATAAGGGAATC |
| | At_MYB29_R | GGCGCCTAGTATAGTTCCC |
| At5g60890.1 | At_MYB34_F | GTGAAGGTGGATGGCGTACT |
| | At_MYB34_R | CCACTTGTTACCCTTAAGAGCATGA |
| At1g18570.1 | At_MYB51_F | CCTCCGTTAACAATCCTCTA *<br>CTACAAGTGTTTCCGTTGACTCTGAA # |

| | | |
|---|---|---|
| | At_MYB51_R | CATCCTCGTGAAGAAACCTC *<br>ACGAAATTATCGCAGTACATTAGAGGA # |
| At2g20610.1 | At_SUR1_F | CGCAAGTTTGATCTTCTTCCCGA |
| | At_SUR1_R | CCGTCTCTGCAACCTTTTTG |
| ***C. rubella / C. bursa-pastoris / C. grandiflora*** | | |
| Carubv10023442m<br>Cagra.18121s0003.1 | Cr_Actin_F | CCGGTATTGTGCTCGACTCT |
| | Cr_Actin_R | AATTTCACGCTCTGCTGTGG |
| Carubv10004524m<br>Cagra.5414s0021.1 | Cr_CYP79B2_F | CGGATCTCATCACTCCCACT |
| | Cr_CYP79B2_R | CTTTAGCATGGCCGGAATCA |
| Carubv10025388m<br>Cagra.7028s0001.1 | Cr_CYP79B3_F | ACCGCCGATGAAATCAAACC |
| | Cr_CYP79B3_R | CCCGACGACTCTGTCTATCT |
| Carubv10007516m<br>Cagra.3807s0022.1 | Cr_CYP83B1_F | GAAACCTCTCCGGCTACCTC |
| | Cr_CYP83B1_R | CTAGCTTTGCCGAGGAGATG |
| Carubv10020723m<br>Cagra.0876s0043.1 | Cr_MYB122_F | GCTGTAGAGCAGAAGGGTTGA |
| | Cr_MYB122_R | TCTCAGTCTGCAGCTTTTGC |
| Carubv10027861m<br>Cagra.0542s0036.1 | Cr_MYB28_F | AGCTCAATGCCTTCTCTGT |
| | Cr_MYB28_R | CAGAAGCCGACAATATATCTTTG |
| Carubv10001328m<br>Cagra.2925s0017.1 | Cr_MYB29_F | ATACGGTGCTTCCTTGACATC |
| | Cr_MYB29_R | GTTGTCCATCTGAAGTTGATCACTA |
| Carubv10027524m<br>Cagra.2519s0060.1 | Cr_MYB34_F | GTGAAGGTGGATGGCGTACT |
| | Cr_MYB34_R | CCACTTGTTACCCTTAAGAGCATGA |
| Carubv10009563m<br>Cagra.4153s0023.1 | Cr_MYB51_F | CCTCCGTTAACAATCCTCCA |
| | Cr_MYB51_R | CGTCCTCGTGAAGAAACCTC |
| ***Cam. sativa*** | | |
| Csa04g051240.1<br>Csa06g040380.1<br>Csa05g016060.1 | Cs_Actin_F | CTTCATGCTGCTTGGTGC |
| | Cs_Actin_R | TTAATTCTCCAGCTATGTATG |
| Csa16g038030.1 | Cs_MYB122_F | CCATTGGAACACTCACATCAA |

| | | |
|---|---|---|
| Csa07g044350.1<br>Csa09g078700.1 | Cs_MYB122_R | GCTACTCTGTTTAAGAACCGA |
| Csa11g097210.1<br>Csa18g034740.1<br>Csa02g068380.1 | Cs_MYB28_F | GAGACAGAGAAGGCATTGA |
| | Cs_MYB28_R | AGAACGGACAACGAGATCAA |
| Csa13g009720.1<br>Csa08g056690.1<br>Csa20g010130.1 | Cs_MYB29_F | ATCATTGAGACATGGAGGAGA |
| | Cs_MYB29_R | GTGATGAGACACCGAGAC |
| Csa17g024370.1<br>Csa14g023900.1<br>Csa03g022440.1 | Cs_MYB51_F | TCCTCCGTTAACAATCATCTA |
| | Cs_MYB51_R | TGTCTTGACGTTCATAGACC |
| **B. Plasmid preparation** | | |
| **prom*AtMYB34* + *AtMYB34*** | | |
| At5g60890.1 | AtMYB34_1_Fw | CGTGCGAUGCCAAGTTTGTACAAAAAAGCAGG |
| | AtMYB34_1_Rv | CACGCGAUTCAGACAAAGACTCCAACCATATTG |
| **prom*AtMYB34*** | | |
| At5g60890.1 | promAtMYB34_Fw | ATATATATATATGGCGCGCCGGTGAGCACAAACTTGTGTT |
| | promAtMYB34_Rv | TTAGCCTGCAGGCTCTCTGCTCTTCTTCTTGA |
| ***CrMYB34*** | | |
| Carubv10027524m | CrMYB34_Fw | ATATAGCCTGCAGGATGGTGAGGACACCATGTTG |
| | CrMYB34_Rv | GCGCGGAATTCGTGTTCTTATTCGTTCCAACAACC |
| **C. Genotyping of *cyp83b1* and *myb* mutant lines** | | |
| ***cyp83b1* SALK_028573** | | |
| At4g31500.1 | *cyp83b1* LP | ATGGTACCGATGGATTCTTCC |
| | *cyp83b1* RP | TCAAGGCCATGATATTGGTTC |
| | SALK LBb1.3 | ATTTTGCCGATTTCGGAAC |
| ***myb34* WiscDsLox424F3** | | |
| At5g60890.1 | *myb34* LP | TCTTCGTTCCAGGAATCAATG |
| | *myb34* RP | AAAGGAGCTTGGACTCCTGAG |
| | Wisc P745 LB | AACGTCCGCAATGTGTTATTAAGTTGTC |
| ***myb51* GK-228B12** | | |

| | | |
|---|---|---|
| At1g18570.1 | *myb51* LP | CAGAGAACGTGGACGAAGAAC |
| | *myb51* RP | TATGATCATACGCCCCATTTC |
| | GABI o8409 LB | ATATTGACCATCATACTCATTGC |
| ***myb122* WiscDsLoxHs206_04H** | | |
| At1g74080.1 | *myb122* LP | CAAGAATGGTACGGACGCCGTGTTG |
| | *myb122*RP | TCCAAAATAATTGTCAATCCCTTCACA |
| | Wisc p745 LB | AACGTCCGCAATGTGTTATTAAGTTGTC |

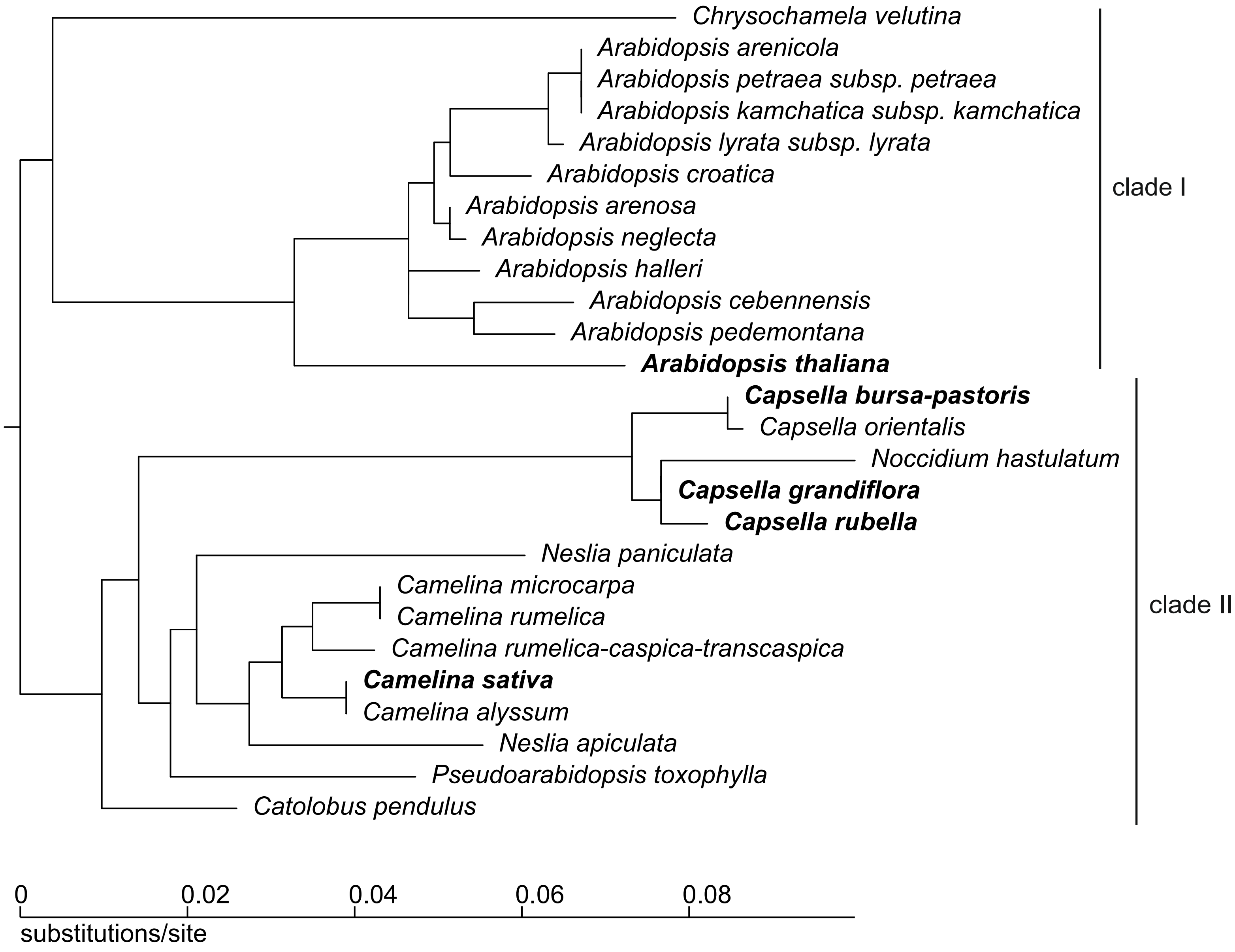


**Fig. S1.** Phylogenetic tree representing relationships between species belonging to the Camelineae tribe. Phylogenetic relations are based on BrassiBase database (Kiefer *et al.*, 2014). Bold letters indicate species tested in this study.

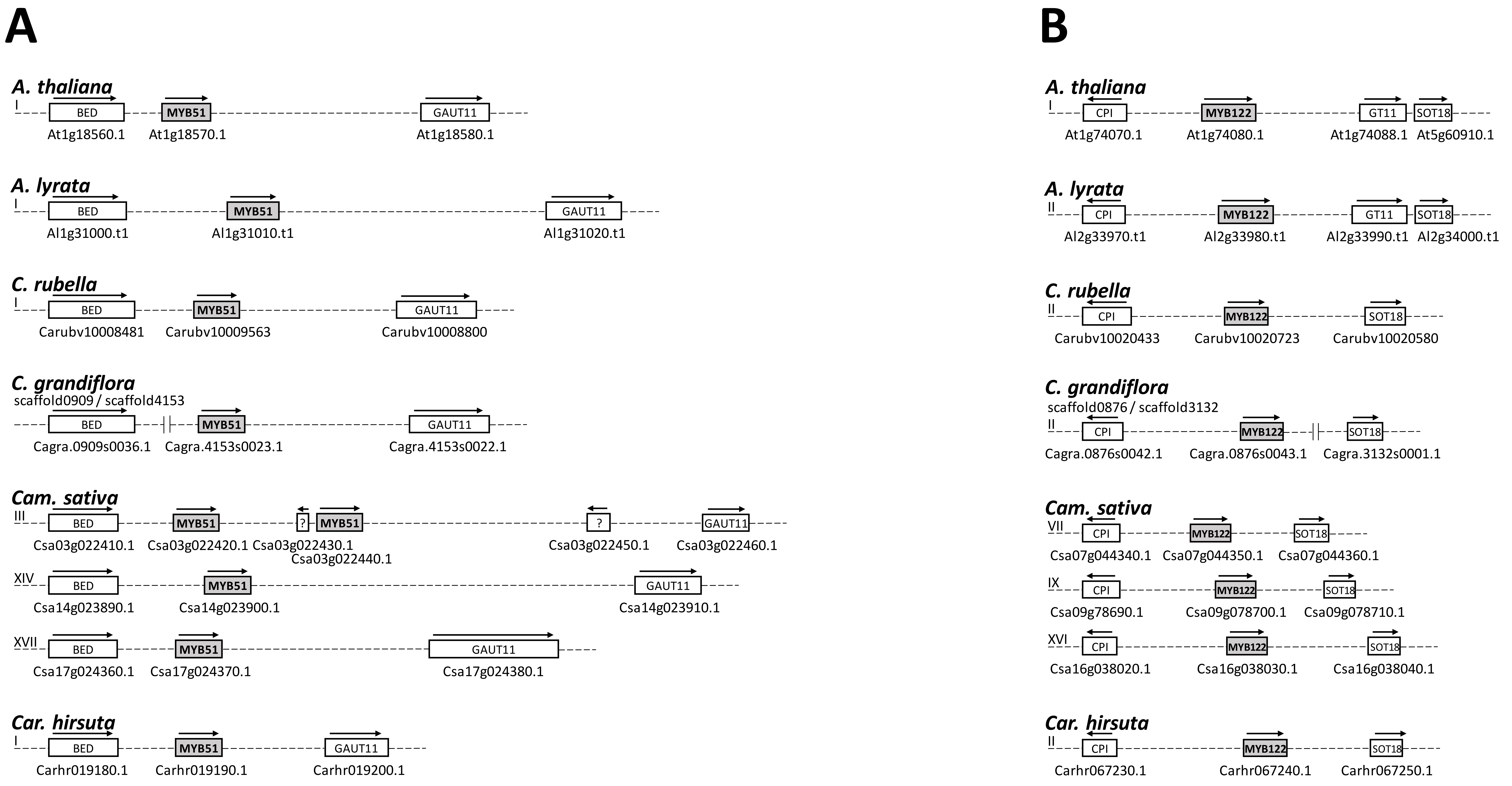


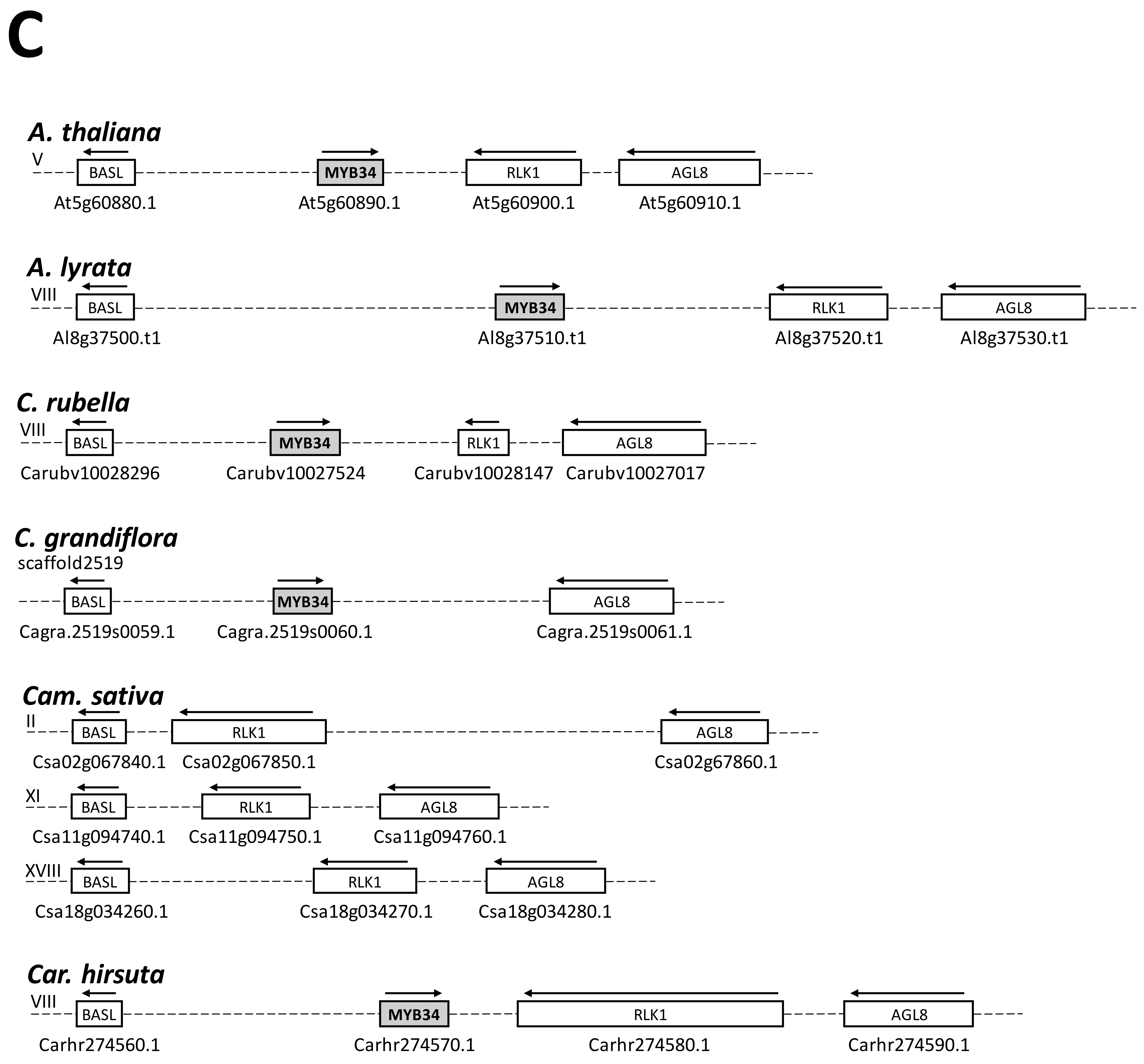


**Fig. S2.** Graphical representation of genomic regions containing *MYB51* (**A**), *MYB122* (**B**), and *MYB34* (**C**) orthologs in the investigated Camelineae species. Roman numerals indicate chromosome numbers (if available). Gene orientations are indicated by arrows. *Cardamine hirsuta* (Cardamineae tribe) has been included for reference.

# A

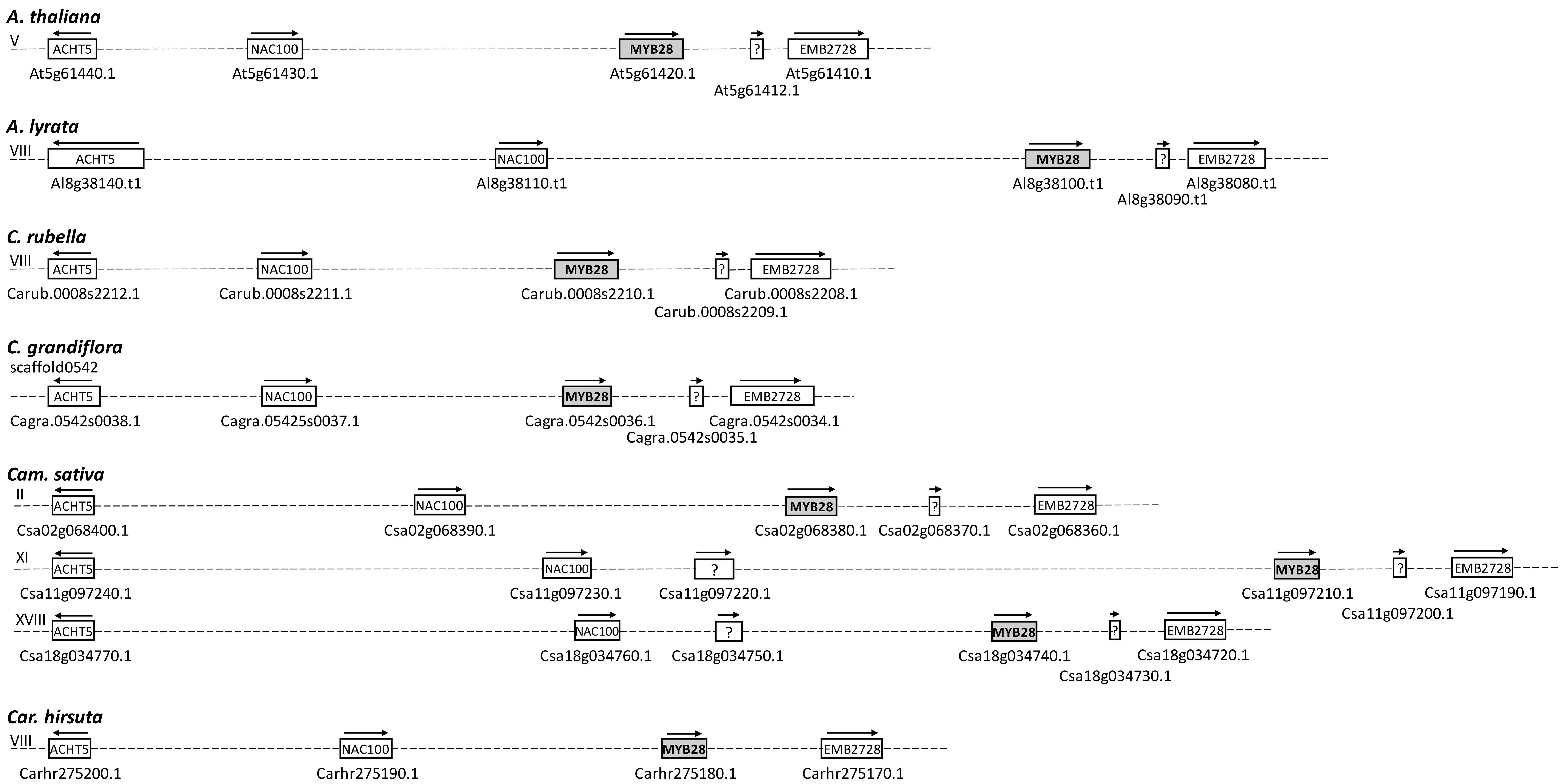


# B

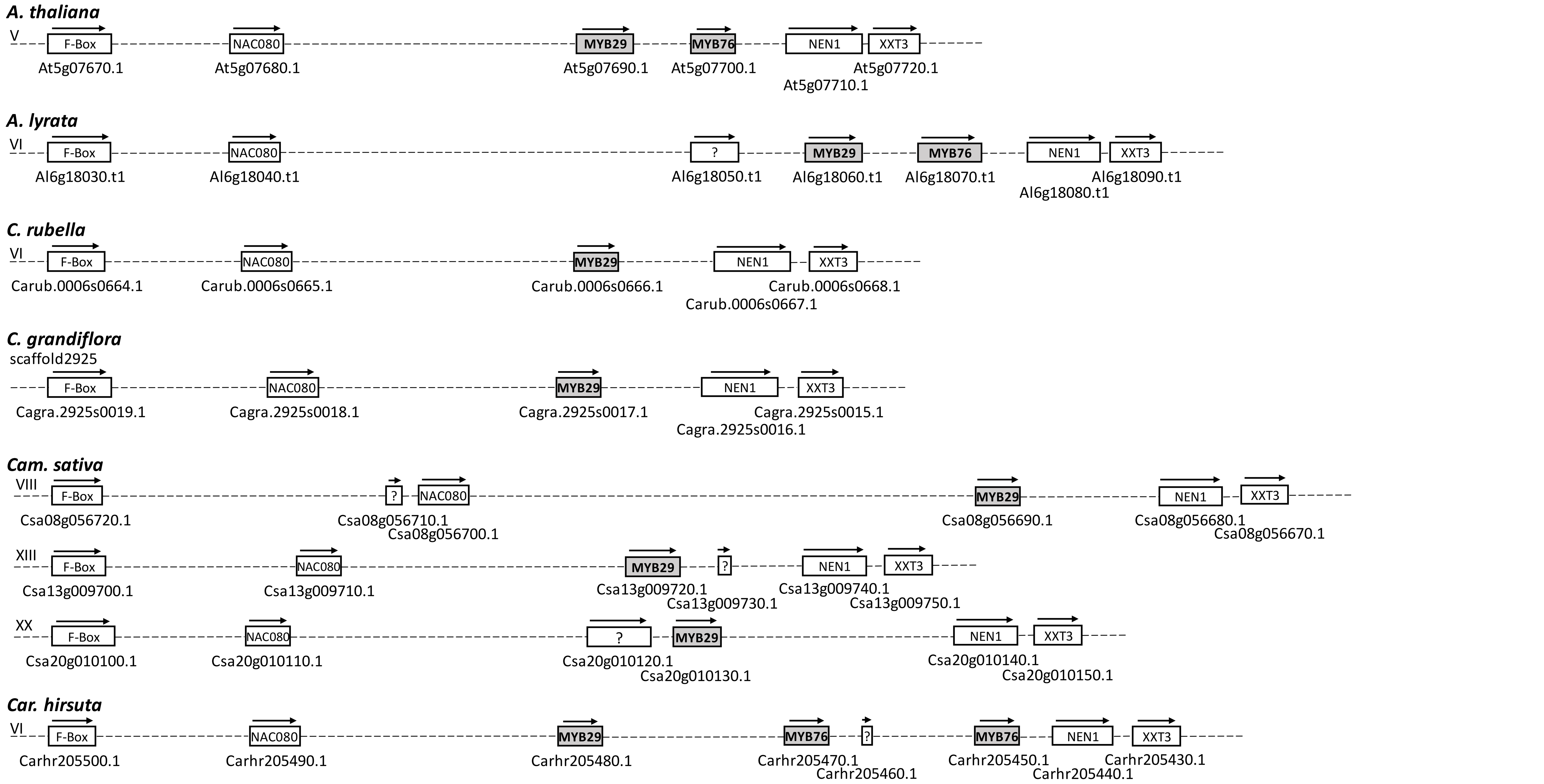


**Fig. S3.** Graphical representation of genomic regions containing *MYB28* (**A**) and *MYB29*/*MYB76* (**B**) orthologs in the investigated Camelineae species. Roman numerals indicate chromosome numbers (if available). Gene orientations are indicated by arrows. *Cardamine hirsuta* (Cardamineae tribe) has been included for reference.

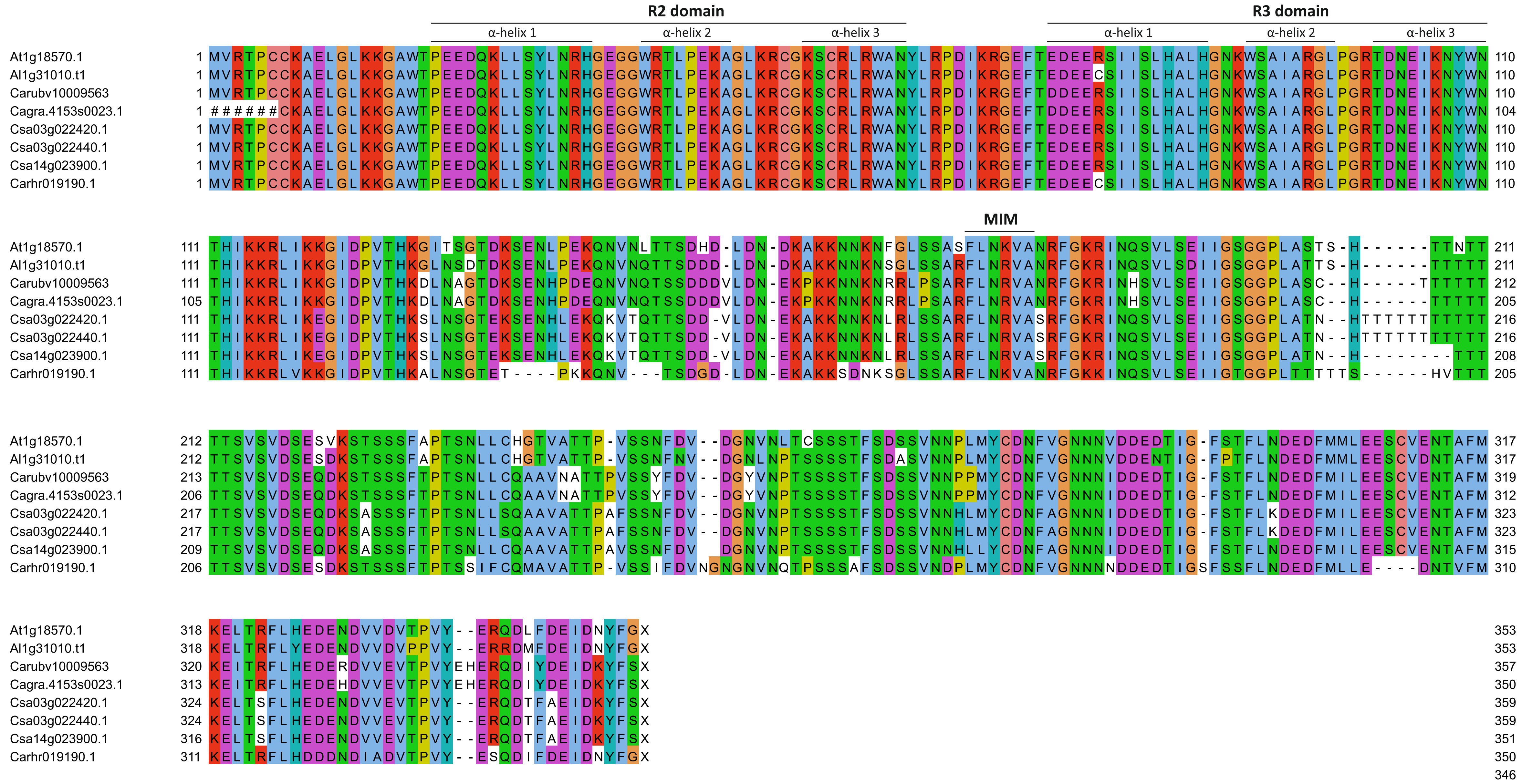


**Fig. S4.** Alignment of proteins encoded by *MYB51* orthologs identified in the investigated Camelineae species. R2, R3 and MIM (MYC-interaction motif) domains are indicated. MYB51 from *Cardamine hirsuta* (Cardamineae tribe) has been included for reference.

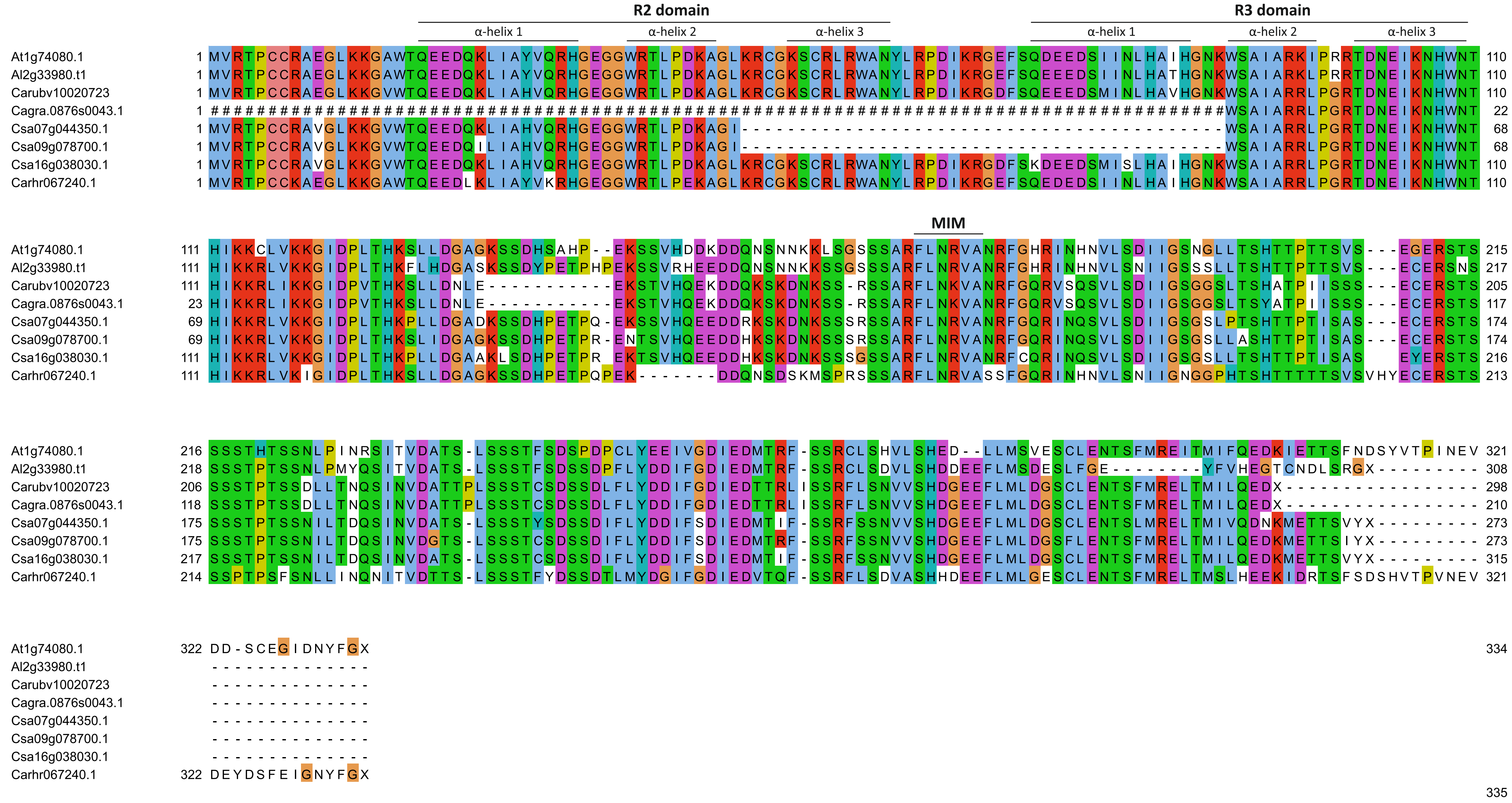


**Fig. S5.** Alignment of proteins encoded by *MYB122* orthologs identified in the investigated Camelineae species. R2, R3 and MIM (MYC-interaction motif) domains are indicated. MYB122 from *Cardamine hirsuta* (Cardamineae tribe) has been included for reference.

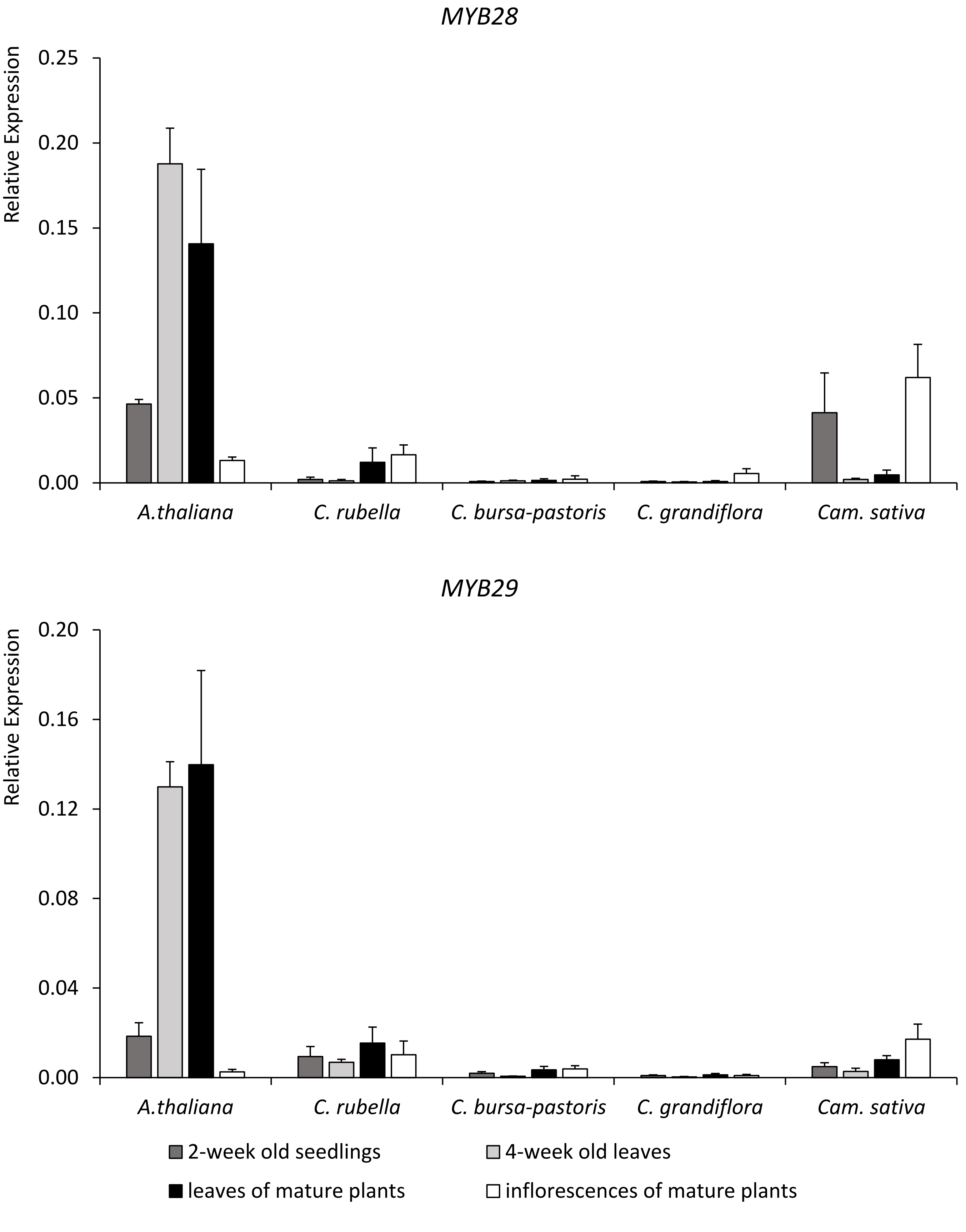


**Fig. S6.** Relative expression levels of *MYB28* and *MYB29* orthologs in investigated Camelineae plants. Results are means ± SD from three independent experiments, each with four biological replicates (n = 12).

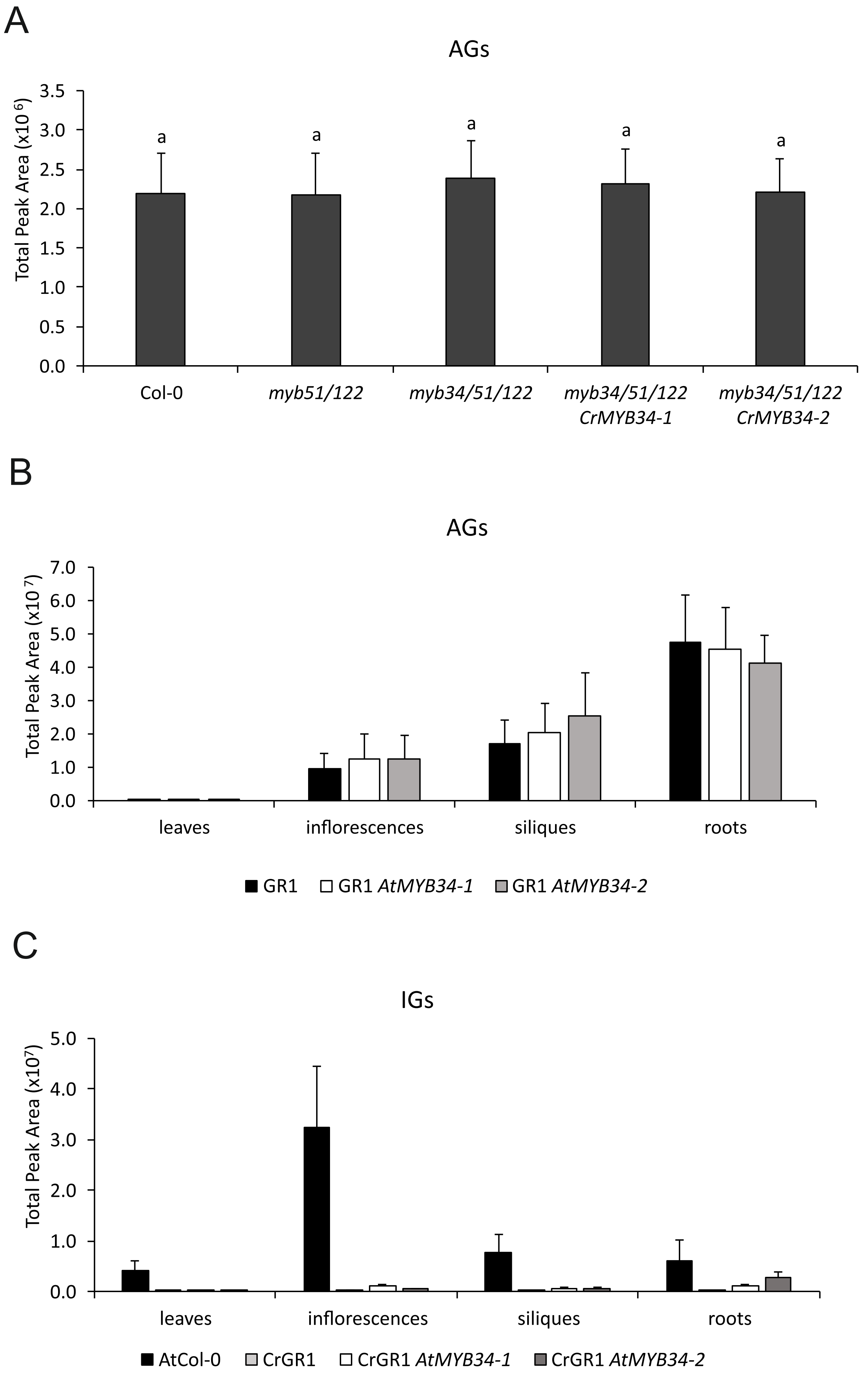


**Fig. S7.** Total accumulation of glucosinolates. (**A**) Aliphatic glucosinolates (AGs) in leaves of *A. thaliana myb* mutant and transgenic lines. (**B**) AGs in different organs of *C. rubella* transgenic plants expressing *AtMYB34*. (**C**) Indolic glucosinolates (IGs) in transgenic *C. rubella* and *A. thaliana* wild-type (Col-0) plants. Bar graphs indicate total peak areas of particular molecular ions corresponding to AGs or IGs detected during LC-MS analysis. Significantly different statistical groups are indicated by ANOVA ($P < 0.05$, Tukey's test). Results are means ± SD from five (A) or eight (B, C) biological replicates.

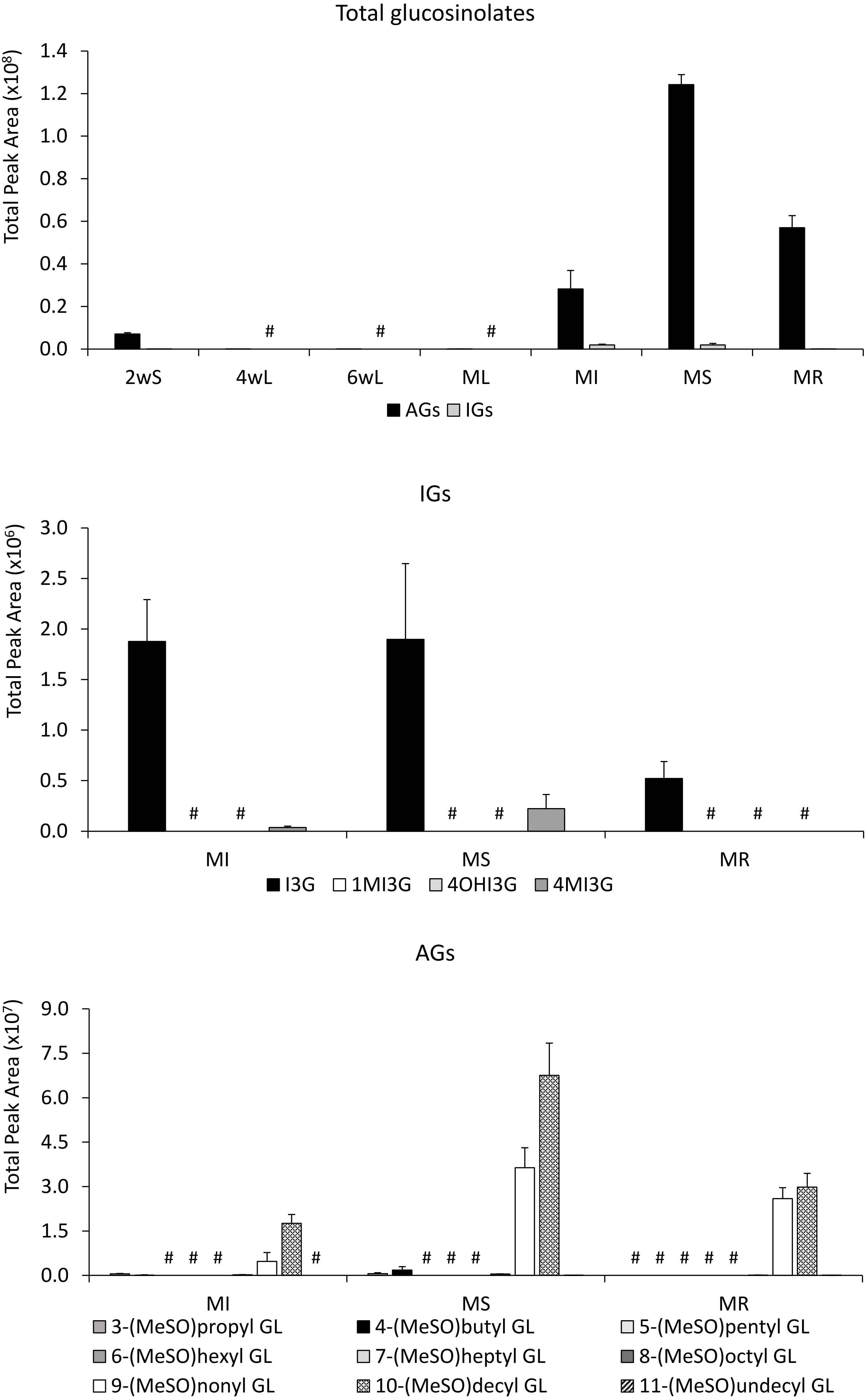


**Fig. S8.** Total accumulation of aliphatic (AGs) and indolic (IGs) glucosinolates in different organs and developmental stages of *C. rubella* GR1 ecotype. Bar graphs indicate peak areas of particular molecular ions detected during LC-MS analysis. Presented values are means ± SD from three independent experiments with three biological replicates in each (n = 9). MeSO – methylsulfinylalkyl, GL – glucosinolate, I3G – 3-indolylmethyl GL, 1MI3G – 1-methoxy-I3G, 4OHI3G – 4-hydroxy-I3G, 4MI3G – 4-methoxy-I3G, 2wS – 2-weeks old seedlings, 4wL – 4-weeks old leaves, 6wL – 6-weeks old leaves, ML – leaves of mature plants, MI – inflorescences of mature plants, MS – siliques of mature plants, MR – roots of mature plants, # - not detected

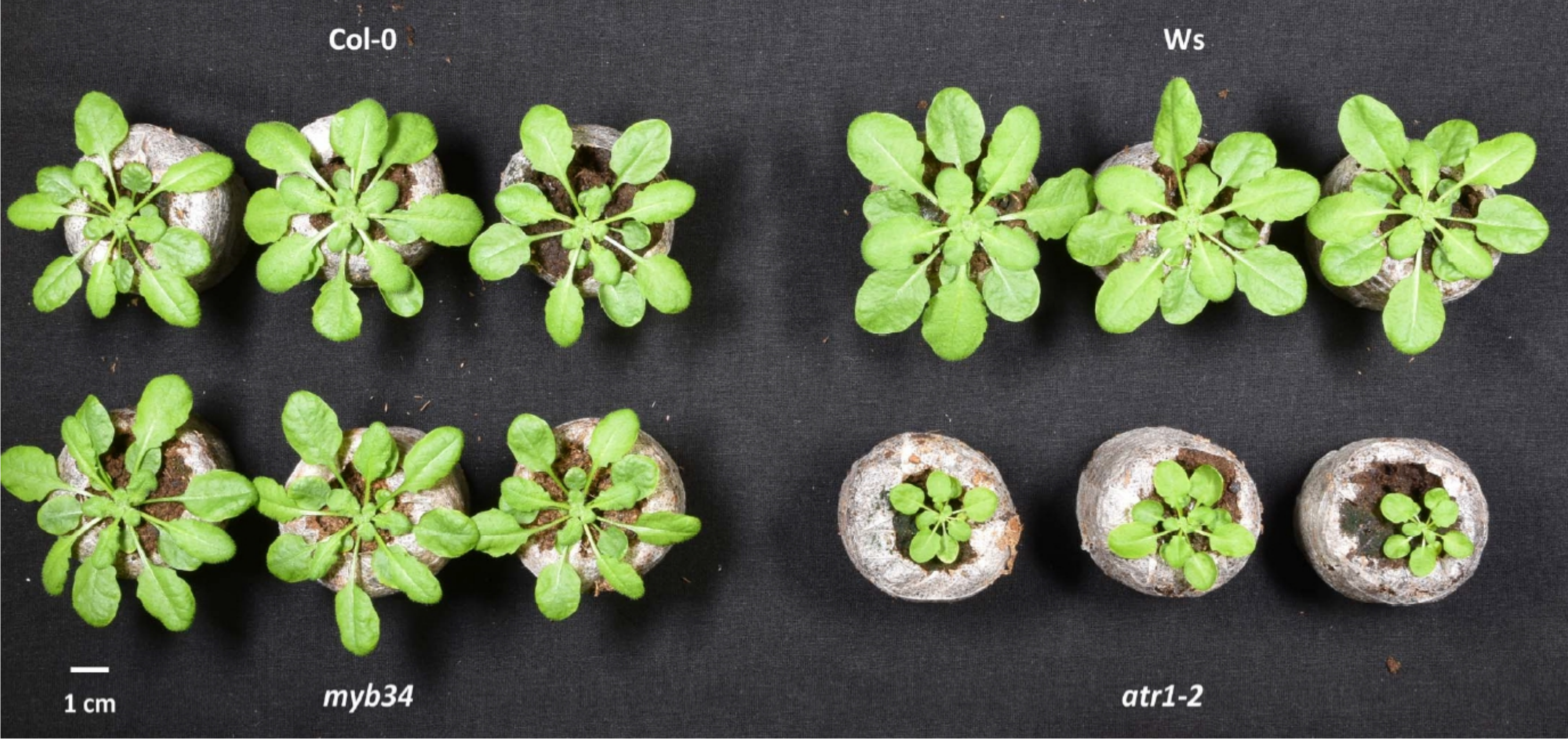


**Fig. S9.** Comparison of growth phenotypes of 6-weeks old Col-0, Ws, *myb34*, and *atr1-2* plants.

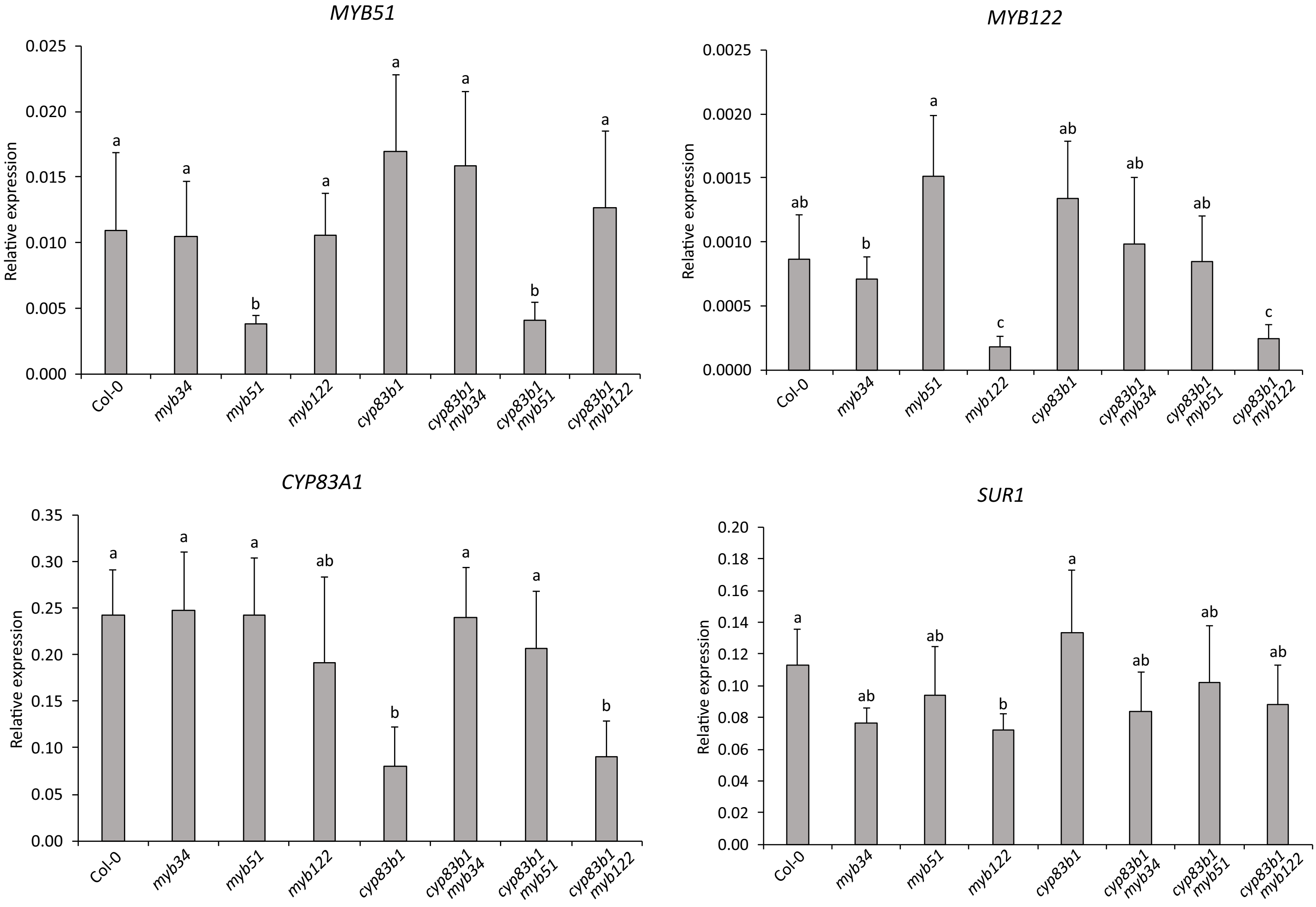


**Fig. S10.** Relative expression levels of *MYB51*, *MYB122*, *CYP83A1*, and *SUR1* genes in *A. thaliana* single and double *cyp83b1* and *myb* mutant lines. Results are means ± SD from two experiments, each with four biological replicates (n = 8). Significantly different statistical groups are indicated based on ANOVA ($P < 0.05$, Games-Howell's test).

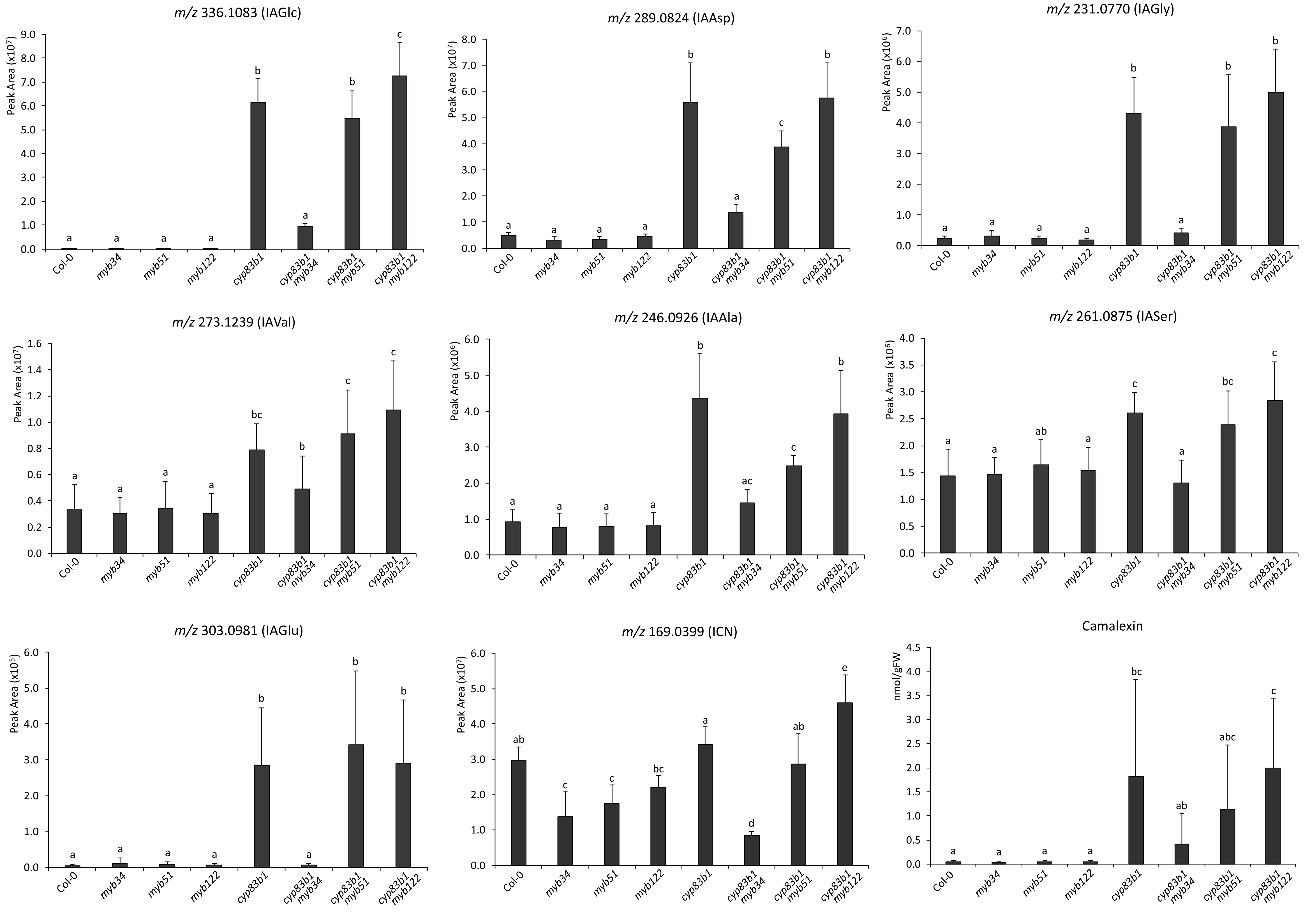


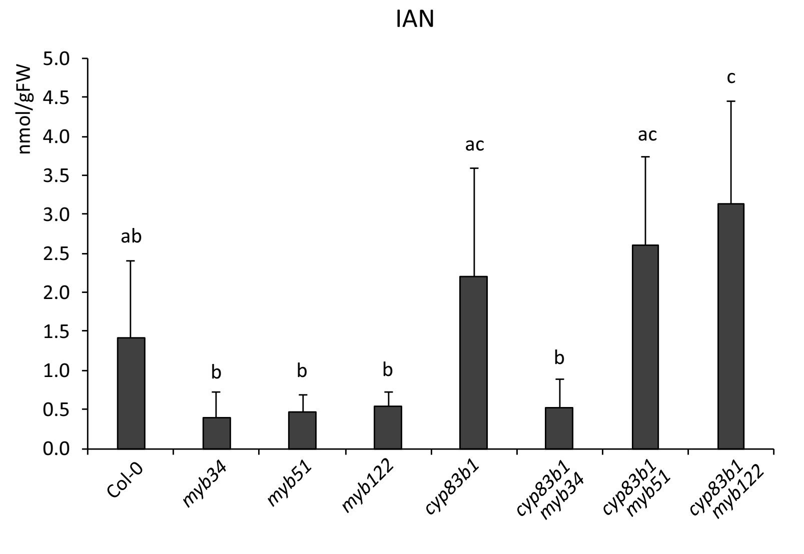


**Fig. S11.** Impact of *cyp83b1* and *myb* mutations on accumulation of different Trp-derivatives in *A. thaliana* leaves. Accumulation levels of putatively identified IAA-conjugates and indol-3-ylmethyl carbonyl nitrile (ICN) presented as peak areas of respective molecular ions detected during LC/MS analysis. Concentrations of camalexin and indole-3-acetonitrile (IAN) calculated based on HPLC-FLD analysis. Results are means ± SD from two experiment, each with four biological replicates (n = 8). Significantly different statistical groups are indicated based on ANOVA ($P < 0.05$, Tukey's test).
